# Metatranscriptomic reanalysis reveals common and specific photosynthetic acclimation of cyanobacteria across diverse natural environments

**DOI:** 10.64898/2026.11.26.769348

**Authors:** Takako Ogawa, Jun Minagawa, Yukako Hihara

## Abstract

Laboratory studies have elucidated acclimation mechanisms in model cyanobacteria, yet their conservation across species and natural environments remains unclear. To address this, we comparatively analyzed publicly available time-series metatranscriptomic datasets from seven distinct natural environments. For each environment, we selected putative dominant species based on read-mapping rates across phylogenetically diverse cyanobacterial genomes. Mapping mRNA reads to the genomes of these putative dominant species revealed both common and environment- or species-specific transcriptional responses. Common responses shared across habitats included a phase shift in diurnal expression of the photosystem II and photosystem I reaction center genes, *psbA* and *psaA*, respectively, and co-induction of *psbA* with genes encoding known high-light acclimation factors, including orange carotenoid protein, high-light-inducible proteins, and SigD, as well as genes belonging to two widely conserved orthogroups of unknown function. Specific responses included daylight-dependent induction of *ndhD1*, encoding a subunit of type 1 NAD(P)H dehydrogenase, in the hot spring cyanobacterium and of *gvp*, encoding gas vesicle structural proteins, in the eutrophic lake cyanobacterium. Furthermore, profiling regulatory genes potentially underlying these acclimation responses, such as those encoding two-component systems and transcription factors, demonstrated that metatranscriptomic data can be used to examine low-abundance regulatory genes as well as highly expressed photosynthetic genes.

## Introduction

Cyanobacteria, the first organisms to perform oxygen-evolving photosynthesis, have changed the composition of the Earth’s atmosphere and expanded their habitats to almost all environments globally. To elucidate the molecular mechanisms that enable cyanobacteria to acclimate flexibly to a wide range of environments, their responses to individual environmental factors have been extensively studied in the laboratory, mainly using genetically tractable model species. These studies have revealed in detail the molecular mechanisms of acclimation responses to changes in environmental factors such as high light (Muramatsu and Hihara 2012), low temperature (Sinetova and Los 2016), high temperature (Rajaram et al. 2014), salt stress (Klähn et al. 2021), iron deficiency (Hernández-Prieto et al. 2012), and carbon/nitrogen balance (Forchhammer and Selim 2020), from the perception of environmental signals to the regulation of gene expression and resulting physiological changes. These studies have shown that protecting photosynthetic complexes from various environmental stresses and balancing the energy supply by photosynthesis with its consumption by cellular metabolism are particularly important for cyanobacteria to survive in fluctuating environments.

However, natural environments differ fundamentally from the laboratory. Rather than a single species experiencing a controlled change of one environmental factor, natural cyanobacterial populations face the diel light cycle superimposed on simultaneous and often unpredictable fluctuations in many other environmental factors, all within diverse microbial communities. Acclimation responses in situ have been examined by qRT-PCR of selected photosynthesis genes, for example, in thermophilic *Synechococcus* in hot spring mats (Jensen et al. 2011) and in marine *Prochlorococcus* (Fernández-Pinos et al. 2015). Metatranscriptomic approaches have further been used to reveal environmental responses of cyanobacteria, such as diel metabolic rhythms of a *Microcystis* bloom (Davenport et al. 2019), thermal acclimation strategies of *Prochlorococcus* (Alonso-Sáez et al. 2023), and acclimation to nitrogen and iron scarcity of marine *Synechococcus* and *Prochlorococcus* (Blaskowski et al. 2025).

Each of these studies, however, has focused on a single habitat type and its resident taxa. Whether the acclimation mechanisms characterized in model strains in the laboratory operate broadly in natural populations, whether some mechanisms are unique to particular environments, and whether there exist acclimation responses that have not been identified in laboratory studies all remain open questions. Here, to address these questions, we comparatively analyzed publicly available time-series metatranscriptome data covering both daytime and nighttime, obtained from diverse environments. Reads were mapped to the genomes of "putative dominant species" selected for each environment from a broad range of cyanobacterial strains, and the expression profiles of photosynthesis-related and regulatory genes were examined.

These analyses revealed responses common among cyanobacteria regardless of their habitats, such as a phase shift in diurnal expression of photosystem II (PSII) and photosystem I (PSI) genes, *psbA* and *psaA*, respectively, and co-induction of *psbA*, several genes related to high-light acclimation, and two widely conserved uncharacterized genes upon daylight exposure. We also found environment- or species-specific responses, such as daylight-dependent induction of a gene encoding a subunit of type 1 NAD(P)H dehydrogenase (NDH-1) complex in the hot spring cyanobacterium and of *gvp* encoding gas vesicle structural proteins in the eutrophic lake cyanobacterium. We also showed that metatranscriptomic data enable analysis of the expression profiles of low-abundance regulatory genes, such as those encoding two-component systems and transcription factors.

## Results and Discussion

We obtained metatranscriptomic datasets from eight different environments (Tables 1 and S1) from the Sequence Read Archive (SRA) of the National Center for Biotechnology Information (NCBI) (Leinonen et al. 2011): the ocean (Zehnpfennig et al. 2022), a eutrophic freshwater lake, an oligotrophic freshwater lake, a humic freshwater lake (Linz et al. 2020), a hot spring (Kawai et al. 2021), a coastal environment (Louyakis et al. 2018), a desert (León-Sobrino et al. 2019), and the bottom of a freshwater lake (Grim et al. 2021). Hereafter, these datasets are referred to as Ocean, Eutrophic Lake, Oligotrophic Lake, Humic Lake, Hot Spring, Coast, Desert, and Lake Floor, respectively. Each dataset was a time-series containing at least one daytime and one nighttime sample.

**Table 1.**
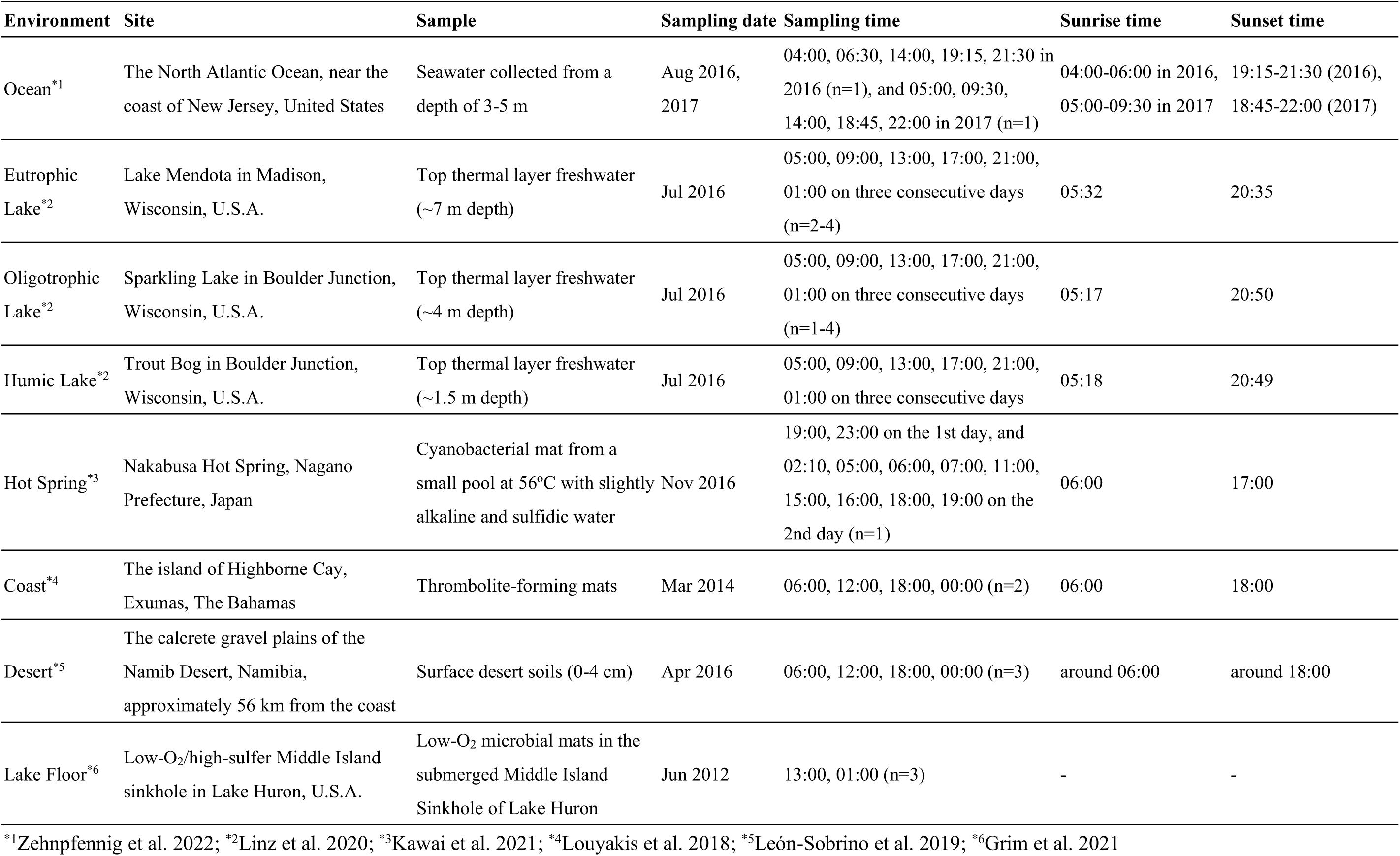
Metatranscriptomic datasets reanalyzed in this study.

| Environment | Site | Sample | Sampling date | Sampling time | Sunrise time | Sunset time |
|---|---|---|---|---|---|---|
| Ocean<sup>*1</sup> | The North Atlantic Ocean, near the coast of New Jersey, United States | Seawater collected from a depth of 3-5 m | Aug 2016, 2017 | 04:00, 06:30, 14:00, 19:15, 21:30 in 2016 (n=1), and 05:00, 09:30, 14:00, 18:45, 22:00 in 2017 (n=1) | 04:00-06:00 in 2016, 05:00-09:30 in 2017 | 19:15-21:30 (2016), 18:45-22:00 (2017) |
| Eutrophic Lake<sup>*2</sup> | Lake Mendota in Madison, Wisconsin, U.S.A. | Top thermal layer freshwater (~7 m depth) | Jul 2016 | 05:00, 09:00, 13:00, 17:00, 21:00, 01:00 on three consecutive days (n=2-4) | 05:32 | 20:35 |
| Oligotrophic Lake<sup>*2</sup> | Sparkling Lake in Boulder Junction, Wisconsin, U.S.A. | Top thermal layer freshwater (~4 m depth) | Jul 2016 | 05:00, 09:00, 13:00, 17:00, 21:00, 01:00 on three consecutive days (n=1-4) | 05:17 | 20:50 |
| Humic Lake<sup>*2</sup> | Trout Bog in Boulder Junction, Wisconsin, U.S.A. | Top thermal layer freshwater (~1.5 m depth) | Jul 2016 | 05:00, 09:00, 13:00, 17:00, 21:00, 01:00 on three consecutive days | 05:18 | 20:49 |
| Hot Spring<sup>*3</sup> | Nakabusa Hot Spring, Nagano Prefecture, Japan | Cyanobacterial mat from a small pool at 56<sup>o</sup>C with slightly alkaline and sulfidic water | Nov 2016 | 19:00, 23:00 on the 1st day, and 02:10, 05:00, 06:00, 07:00, 11:00, 15:00, 16:00, 18:00, 19:00 on the 2nd day (n=1) | 06:00 | 17:00 |
| Coast<sup>*4</sup> | The island of Highborne Cay, Exumas, The Bahamas | Thrombolite-forming mats | Mar 2014 | 06:00, 12:00, 18:00, 00:00 (n=2) | 06:00 | 18:00 |
| Desert<sup>*5</sup> | The calcrete gravel plains of the Namib Desert, Namibia, approximately 56 km from the coast | Surface desert soils (0-4 cm) | Apr 2016 | 06:00, 12:00, 18:00, 00:00 (n=3) | around 06:00 | around 18:00 |
| Lake Floor<sup>*6</sup> | Low-O<sub>2</sub>/high-sulfer Middle Island sinkhole in Lake Huron, U.S.A. | Low-O<sub>2</sub> microbial mats in the submerged Middle Island Sinkhole of Lake Huron | Jun 2012 | 13:00, 01:00 (n=3) | - | - |

<sup>\*1</sup>Zehnpfennig et al. 2022; <sup>\*2</sup>Linz et al. 2020; <sup>\*3</sup>Kawai et al. 2021; <sup>\*4</sup>Louyakis et al. 2018; <sup>\*5</sup>León-Sobrino et al. 2019; <sup>\*6</sup>Grim et al. 2021

### 1. Identification of putative dominant cyanobacterial species in each environment

To map the mRNA reads from each metatranscriptomic dataset in a consistent manner, using the genomic sequence of the dominant cyanobacterial species in each environment as a reference would be ideal. However, some datasets lacked the information required to identify the dominant species, such as metagenomic data or 16S rRNA gene sequences. Thus, we adopted a strategy to define a "putative dominant species" for each dataset by selecting the genome that exhibited the highest mapping rate among the 159 cyanobacterial genomes (Table S2) registered in the Kyoto Encyclopedia of Genes and Genomes (KEGG) database (Kanehisa et al. 2025; https://www.genome.jp/kegg/). These 159 genomes represented a broad range of phylogenetic lineages (Fig. 1) derived from diverse habitats. mRNA reads from the Ocean, Eutrophic Lake, Oligotrophic Lake, and Hot Spring datasets mapped to less than 40% of the 159 genomes, showing a pronounced concentration within a few distinct cyanobacterial clades that differed depending on the environmental conditions (Fig. 1). In contrast, mRNA reads from the Coast, Desert, and Lake Floor datasets mapped broadly across diverse lineages throughout the phylogenetic tree, with a distinct genome displaying a notably high mapping rate in each environment (Fig. 1). For the Humic Lake dataset, no reads mapped to any of the 159 genomes (0% mapping rate; Table S2).

**Figure 1.**
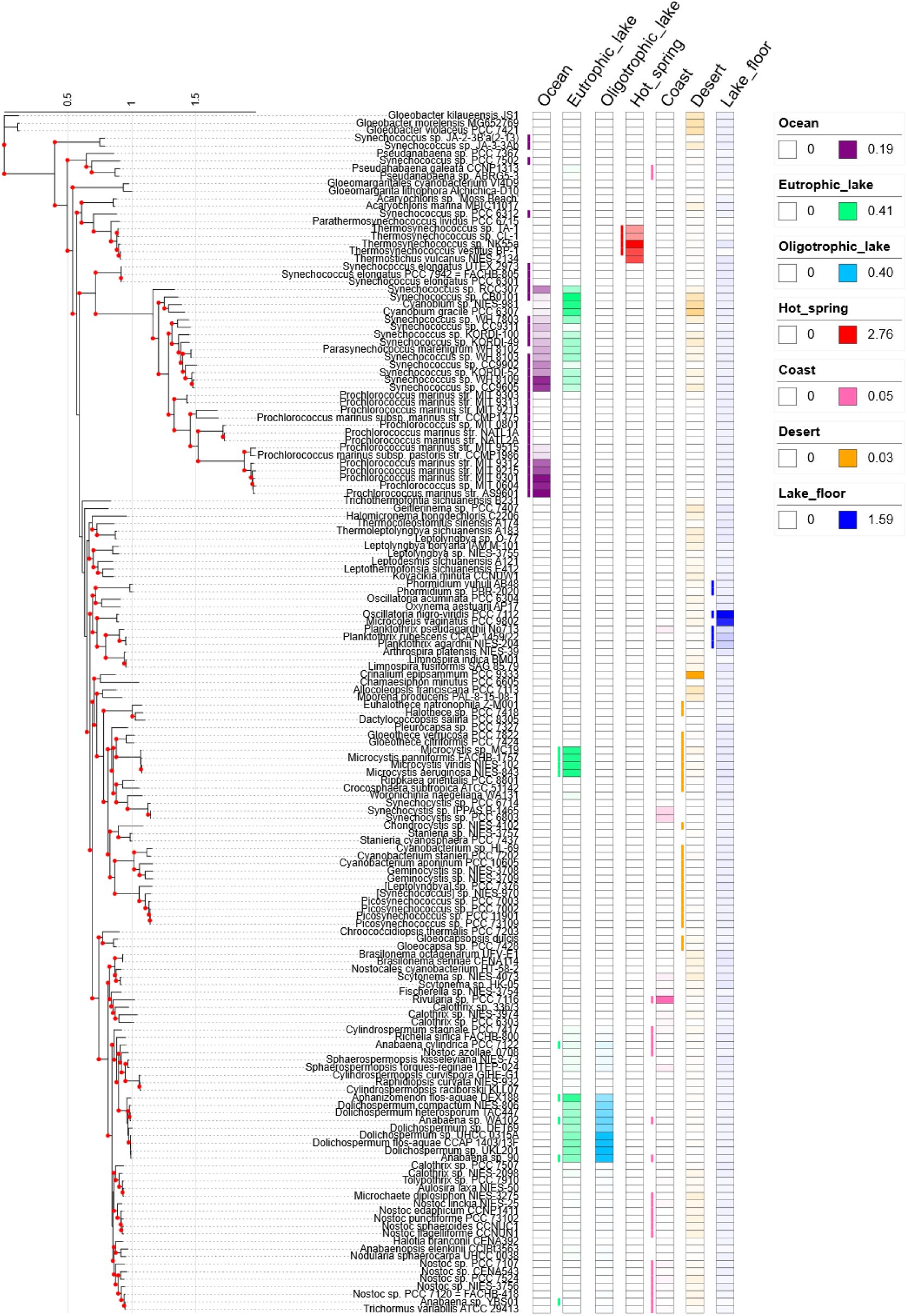
Phylogenetic tree of 159 cyanobacterial strains and heatmaps showing the mapping rates of mRNA reads their genomes. The phylogenetic tree was inferred using the maximum-likelihood method based on a concatenated alignment of 126 single-copy orthologs. The scale bar represents evolutionary distance in amino acid substitutions per site. Red dots indicate nodes with ultrafast bootstrap support values of ≥95. Mapping rates were calculated as the percentage of total mRNA reads in each sample that mapped to each genome. For each environment, the heatmap shows the mean mapping rate across samples marked with an asterisk in Table S1. A separate color scale was used for each environment, with white indicating 0% and the darkest color indicating the highest mapping rate within that environment; intermediate values are represented by a gradient. The bars to the left of each environment’s heatmap indicate the position on the phylogenetic tree of cyanobacterial taxa previously reported at the corresponding sampling sites (Table S4). Taxa are represented at the genus level for the Ocean, Eutrophic Lake, Hot Spring, and Lake Floor datasets, at the family level for the Coast dataset, and at the order level for the Desert dataset.

Cyanobacteria whose genomes showed high mapping rates (>50% of the maximum mapping rate for each dataset; Table S3) in the Ocean, Hot Spring, Coast and Lake Floor datasets were consistent, at the family, genus, or species level, with cyanobacteria previously reported from these sampling sites (Table S4; Fig. 1). In the Eutrophic Lake dataset, two of the three lineages in which genomes with high mapping rates were concentrated corresponded to genera previously reported at the sampling site (Table S4; Fig. 1). Although cyanobacteria belonging to the third lineage (*Synechococcus*/*Cyanobium*) have not been reported from this location, this lineage is widely distributed across both freshwater and marine environments (Wang et al. 2022), making its presence in the eutrophic freshwater lake plausible. Similarly, in the Oligotrophic Lake and Desert datasets, high mapping rates were observed for lineages or species not previously confirmed at these sampling sites (Table S3 and S4). However, these corresponded to freshwater genera (*Anabaena*, *Aphanizomenon* and *Dolichospermum*) and to *Crinalium epipsammum* PCC 9333 (C.PCC9333), an isolate from sandy crust (Zhang et al. 2015), respectively; thus, the presence of related taxa in oligotrophic freshwater or desert environments is reasonable. For the Humic Lake dataset, although cyanobacterial reads have been reported in this sample (Linz et al. 2020), the mapping rate was 0% across all 159 genomes (Table S2). This lack of mapping likely reflects a low abundance of cyanobacteria or the absence of closely related species among the 159 genomes.

From these results, we designated 10 cyanobacterial species as putative dominant species across the seven environmental datasets (Table 2). For the Ocean and Eutrophic Lake datasets, in which genomes with high mapping rates were concentrated in several distinct lineages, we selected the single strain with the highest mapping rate from each lineage. For the other environmental datasets, we selected a single strain showing the highest mapping rate overall. Hereafter, for simplicity, gene expression in each dataset is referred to as expression in these putative dominant species.

**Table 2.** Putative dominant cyanobacterial species selected for each environment.

| **Environment** | **Order** | **Family** | **Species** | **(abbreviation)** | **Mapping rate (%)** |
|---|---|---|---|---|---|
| **Ocean** | Synechococcales | Synechococcaceae | *Synechococcus* sp. WH 8109 | (S.WH8109) | 0.163 |
|  | Synechococcales | Prochlorococcaceae | *Prochlorococcus marinus* AS9601 | (P.AS9601) | 0.191 |
| **Eutrophic Lake** | Synechococcales | Synechococcaceae | *Synechococcus* sp. CB0101 | (S.CB0101) | 0.406 |
|  | Chroococcales | Microcystaceae | *Microcystis aeruginosa* NIES-843 | (M.NIES-843) | 0.368 |
|  | Nostocales | Aphanizomenonaceae | *Aphanizomenon flos-aquae* DEX188 | (A.DEX188) | 0.325 |
| **Oligotrophic Lake** | Nostocales | Aphanizomenonaceae | *Dolichospermum* sp. UHCC 0315A | (D.UHCC0315A) | 0.386 |
| **Humic Lake** | - | - | - |  | - |
| **Hot Spring** | Acaryochloridales | Thermosynechococcaceae | *Thermosynechococcus* sp. NK55 | (T.NK55) | 2.764 |
| **Coast** | Nostocales | Rivulariaceae | *Rivularia* sp. PCC 7116 | (R.PCC 7116) | 0.050 |
| **Desert** | Gomontiellales | Gomontiellaceae | *Crinalium epipsammum* PCC 9333 | (C.PCC 9333) | 0.029 |
| **Lake Floor** | Oscillatoriales | Oscillatoriaceae | *Oscillatoria nigro-viridis* PCC 7112 | (O.PCC 7112) | 1.588 |

### 2. Proportion of genes with detectable expression and functional categories of highly expressed genes

In each putative dominant species, the proportion of genome-encoded genes with detectable expression [Transcripts Per Million (TPM) > 0; hereafter, "detected genes"] tended to increase with the number of mRNA reads mapped to the genome (Fig. S1). This proportion varied widely among the putative dominant species, exceeding 90% in *Thermosynechococcus* sp. NK55 (T.NK55) for the Hot Spring dataset, whereas it was only approximately 1% in *Rivularia* sp. PCC 7116 (R.PCC7116) for the Coast dataset (Figs. S1 and S2). On the other hand, in cases such as the three putative dominant species for the Eutrophic Lake dataset, the proportion of detected genes differed despite similar numbers of mapped mRNA reads (Fig. S1). This difference might be attributed to varying genetic distances between the actual dominant species and the putative dominant species. In the putative dominant species with greater genetic divergence from the actual dominant species, mRNA reads might have mapped preferentially to gene sequences that are highly conserved across species, thereby decreasing the overall proportion of detected genes.

To characterize the functional categories of the detected genes in each putative dominant species, we examined the proportion of the detected genes in each category of the KEGG PATHWAY database. In *Microcystis aeruginosa* NIES-843 (M.NIES-843), *Aphanizomenon flos-aquae* DEX188 (A.DEX188), *Dolichospermum* sp. UHCC 0315A (D.UHCC0315A), and T.NK55, the proportion of detected genes exceeded 80% across most categories, whereas it remained below 50% in *Synechococcus* sp. WH 8109 (S.WH8109), *Prochlorococcus marinus* AS9601 (P.AS9601), *Synechococcus* sp. CB0101 (S.CB0101), R.PCC7116, C.PCC9333, and *Oscillatoria nigro-viridis* PCC 7112 (O.PCC7112) (Figs. 2A and S2). In "Energy metabolism", the proportion of detected genes exceeded 50% in seven of the ten putative dominant species (Fig. 2A). This might reflect the high sequence conservation and/or high expression levels of genes assigned to this category. The proportion of detected genes in "Transcription" also exceeded 50% even in the six species that had low overall proportions of detected genes (Fig. 2A). This might be attributed to the small number of genes assigned to this category (n=5).

**Figure 2.**
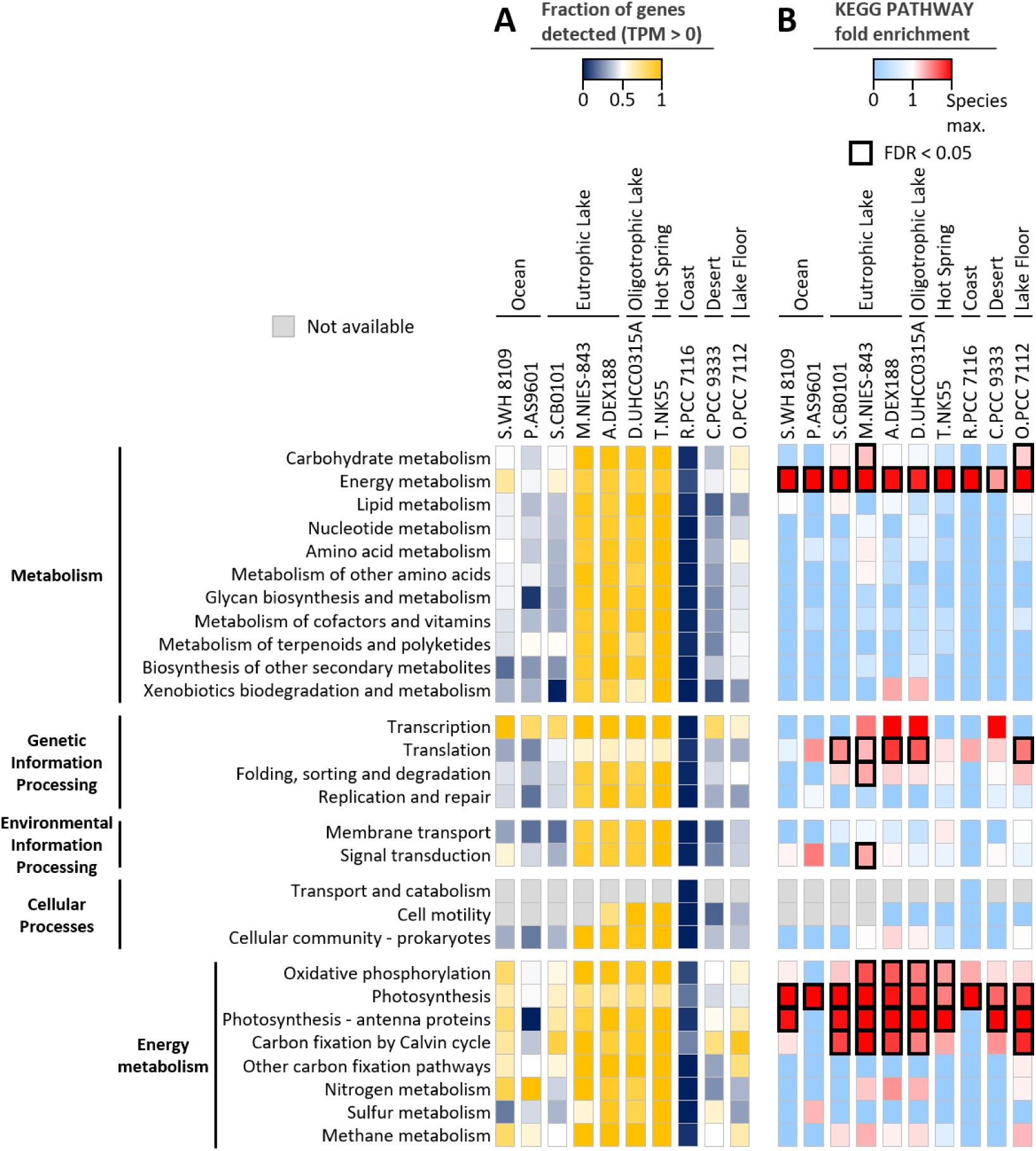
Heatmaps showing (A) the proportion of genes detected (TPM > 0) and (B) fold enrichment of highly expressed genes in each KEGG PATHWAY category. For each putative dominant species, the sample collected at the time point closest to noon (daytime sample) is presented. In both A and B panels, categories containing no genes are shown in light gray. Fold enrichment was calculated for highly expressed genes, defined as the top 10% of the detected genes ranked by TPM in each sample.

To investigate which functional categories were highly expressed in each putative dominant species, we performed KEGG PATHWAY enrichment analysis focusing on the top 10% of detected genes ranked by TPM, hereafter referred to as "highly expressed genes" (Figs. 2B and S3). In R.PCC7116, the number of detected genes was limited to only 35 to 71 across all time points (Fig. S2), resulting in very few highly expressed genes. Therefore, the results of the enrichment analysis for this species had low statistical reliability and should be regarded as indicative only. Across all putative dominant species, "Energy metabolism" tended to be enriched (fold enrichment > 1). In contrast, genes assigned to "Translation" were significantly enriched among the highly expressed genes [false discovery rate (FDR) < 0.05]. only in the putative dominant species for the Eutrophic Lake, Oligotrophic Lake, and Lake Floor datasets. In M.NIES-843 and O.PCC7112, several KEGG PATHWAY categories, such as "Carbohydrate metabolism", "Signal transduction", and "Amino acid metabolism", were significantly enriched, whereas these categories were not enriched in the other putative dominant species. While these findings may indicate species-specific high expression of genes related to these pathways, they could also be an artifact of the larger total number of genes encoded in the genomes of M.NIES-843 and O.PCC7112 compared to the other putative dominant species (Fig. S2), which leads to overestimation of fold enrichment values (see Materials and Methods). "Energy metabolism" was enriched among the highly expressed genes in all putative dominant species, suggesting that genes assigned to this category were robustly and highly expressed across all species, irrespective of the gene number-bias.

Among the subcategories of "Energy metabolism", "Photosynthesis" tended to be enriched across all putative dominant species (Figs. 2B and S3). Furthermore, in the sample collected at the time point closest to noon for each dataset (hereafter, "daytime sample"), "Photosynthesis - antenna proteins" was significantly enriched in eight of the ten putative dominant species, whereas "Carbon fixation by Calvin cycle" was significantly enriched in the putative dominant species for the Eutrophic Lake, Oligotrophic Lake, and Lake Floor datasets (Figs. 2B and S3). These results demonstrate that the enrichment of "Energy metabolism" among the highly expressed genes commonly observed across the putative dominant species is primarily driven by the robust expression of genes involved in photosynthesis and related processes. In addition, "Oxidative phosphorylation" was significantly enriched at all time points in M.NIES-843, A.DEX188, and D.UHCC0315A, whereas in T.NK55, it was significantly enriched only in the daytime sample (Fig. S3), suggesting that high expression of genes in the category also contributed to the overall enrichment of "Energy metabolism" within the highly expressed gene sets.

### 3. Expression profiles of genes encoding subunits of photosynthesis-related complexes

Among the genes assigned to "Photosynthesis" in the KEGG PATHWAY database, either *psbA* or *psbD*, which encode the PSII reaction center proteins D1 and D2, respectively, was the most highly expressed gene in daytime samples in all putative dominant species except for O.PCC7112 (Table S5), showing markedly higher TPM values than genes encoding other PSII subunits (Fig. 3A). Because PSII reaction center proteins are susceptible to light-induced damage and the D1 protein in particular is known to undergo rapid turnover (Su et al. 2023), the high expression of these reaction center genes likely reflects the high cellular demand for the D1 and D2 proteins to cope with the continuous PSII photodamage under sunlight. In contrast, *psbH* showed higher TPM values than *psbA* and *psbD* in O.PCC7112 for the Lake Floor dataset (Fig. 3A). In the low-light environment expected at the lake bottom, O.PCC7112 might have a relatively lower need to cope with PSII photodamage. Instead, there might be a greater requirement for PsbH, which contributes to the stabilization of the PSII dimer (Iwai et al. 2006).

**Figure 3.**
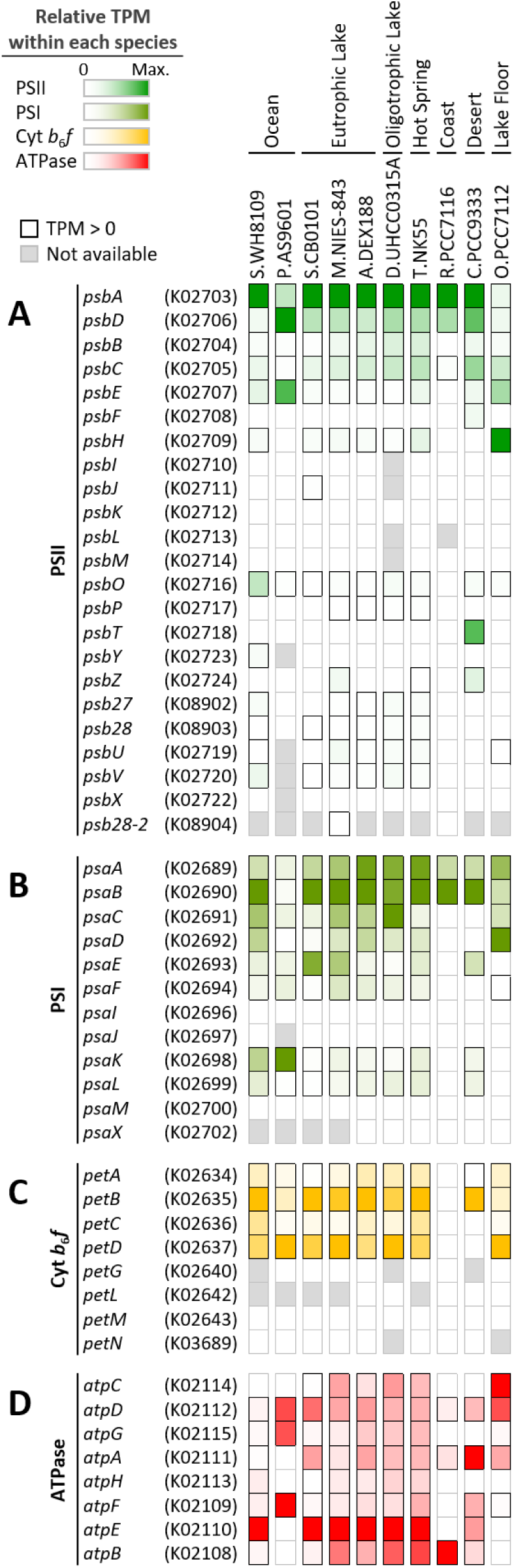
Heatmaps showing the expression profiles of genes assigned to "Photosynthesis" in the KEGG PATHWAY database in the daytime samples of the putative dominant species. (A) Photosystem II (PSII); (B) photosystem I (PSI); (C) cytochrome *b*_6_*f* (Cyt *b*_6_*f*) complex; and (D) ATP synthase (ATPase). When multiple genes were assigned the same KO identifier, their mean TPM was used.

In the case of PSI, several genes encoding subunits critical for electron transfer, such as those involved in iron-sulfur cluster binding or ferredoxin docking (Chen et al. 2022), exhibited expression levels comparable to those of the reaction center genes, *psaA* and *psaB*, in some species (Fig. 3B). These included *psaE* in S.CB0101, *psaC* in D.UHCC0315A, and *psaD* in O.PCC7112. In P.AS9601, the TPM values of *psaK* encoding a small subunit were markedly higher than those of *psaA*, *psaB*, and the genes encoding other major PSI subunits involved in electron transfer (Fig. 3B). In *Synechocystis* sp. PCC 6803 possessing two copies of *psaK*, *psaK2* expression was induced under high-light conditions and PsaK1 in the PSI complex is replaced by PsaK2 under these conditions, suggesting a role of PsaK2 in high-light acclimation (Fujimori et al. 2005). However, the genome of P.AS9601 encodes only a single copy of *psaK*, and whether this gene is involved in high-light acclimation remains unknown.

Regarding the cytochrome *b*_6_*f* (Cyt *b*_6_*f*) complex, the four large subunits responsible for organizing the electron transfer chain are encoded by *petA*, *petB*, *petC*, and *petD* (encoding cytochrome *f*, cytochrome *b*_6_, Rieske iron-sulfur subunit, and subunit IV, respectively; Malone et al. 2019). Although none of these genes showed detectable expression in R.PCC7116 and only a subset showed detectable expression in C.PCC9333, the other putative dominant species shared a characteristic expression profile: *petB* and *petD* were expressed at approximately 2.4- to 260-fold higher levels than *petA* and *petC* (Fig. 3C). This shared expression profile likely reflects the genomic and biogenetic architecture in cyanobacteria, where *petBD* and *petAC* form distinct operons (Schneider et al. 2000), and the initial step of assembly of the Cyt *b*_6_*f* complex involves the transcriptional activation of the *petBD* operon followed by the insertion of its products into the thylakoid membrane (Hasan et al. 2013). A similar pattern of higher expression of *petBD* than *petAC* has also been reported in a metatranscriptomic analysis of *Microcystis* blooms in a lake (Davenport et al. 2019). In all putative dominant species, the TPM values of *petG*, *petL*, *petM*, and *petN*, which encode small subunits and are each shorter than 150 nt, were zero (Fig. 3C), likely reflecting the technical detection limit for short genes.

Across all putative dominant species in which expression of ATP synthase genes was detected, *atpE*, which encodes the c subunit, consistently exhibited the highest TPM, being approximately 3.0- to 6.6-fold higher than the average TPM of the other subunit genes (Fig. 3D). In the Fo sector of cyanobacterial ATP synthase, 13 to 15 or more c subunits assemble into a c-ring (Pogoryelov et al. 2007). Thus, the elevated expression of *atpE* likely reflects the higher stoichiometric demand for c subunits relative to the other subunits.

Taken together, these results indicate a common trend in expression profiles of genes encoding photosynthetic complexes across putative dominant species (Fig. 3). Namely, subunit genes that are involved in electron transfer or are required in higher stoichiometric ratios tended to exhibit relatively high expression levels within their respective complexes. On the other hand, a species-specific expression pattern was observed in photosystem genes, where small subunit genes appeared to be more highly expressed than the reaction center genes (Figs. 3A and 3B). In *Prochlorococcus marinus* str. MIT 9301, the PSI reaction center genes *psaA* and *psaB* are expressed at higher levels than the small subunit genes *psaF* and *psaJ* under laboratory culture conditions. However, when reads from the *Tara* Oceans metatranscriptomic dataset (Salazar et al. 2019) were mapped to the genome of this strain, *psaF* and *psaJ* showed expression levels comparable to *psaA* and *psaB* (Alonso-Sáez et al. 2023). In natural environments, elevated expression of genes encoding small subunits may help cyanobacteria acclimate to local environmental conditions.

### 4. Expression profiles of genes encoding subunits of light-harvesting antenna complexes

All putative dominant species except P.AS9601 possess phycobilisomes as light-harvesting antenna complexes, which exhibit a hemidiscoidal structure with rods radiating from a central core (Bryant and Gisriel 2024). Among the phycobiliproteins binding various bilin chromophores, allophycocyanin (APC) forms the core, whereas phycocyanin (PC) constitutes the rods. In some species, the rods further extend with phycoerythrocyanin (PEC) or phycoerythrin (PE) attached to the distal ends of PC (Stadnichuk et al. 2015). In R.PCC7116, which had the lowest proportion of detected genes relative to the total genome-encoded genes (Fig. S2), expression of most phycobilisome-related genes was undetectable (Fig. 4A). In the remaining eight species, however, many genes encoding phycobiliproteins were included among the highly expressed genes (Table S5) and exhibited high TPM values in daytime samples (Fig. 4A). By comparison, only a subset of genes encoding phycobilisome linker proteins, which connect phycobiliproteins, was included among the highly expressed genes. Thus, a common trend across these putative dominant species was that phycobiliprotein genes tended to have higher TPM values than linker protein genes. For example, in each putative dominant species, the average TPM of *cpcA* and *cpcB*, which encode PC, was approximately 5–21-fold higher than that of *cpcC*, which encodes a PC-associated rod linker protein (Fig. 4A).

**Figure 4.**
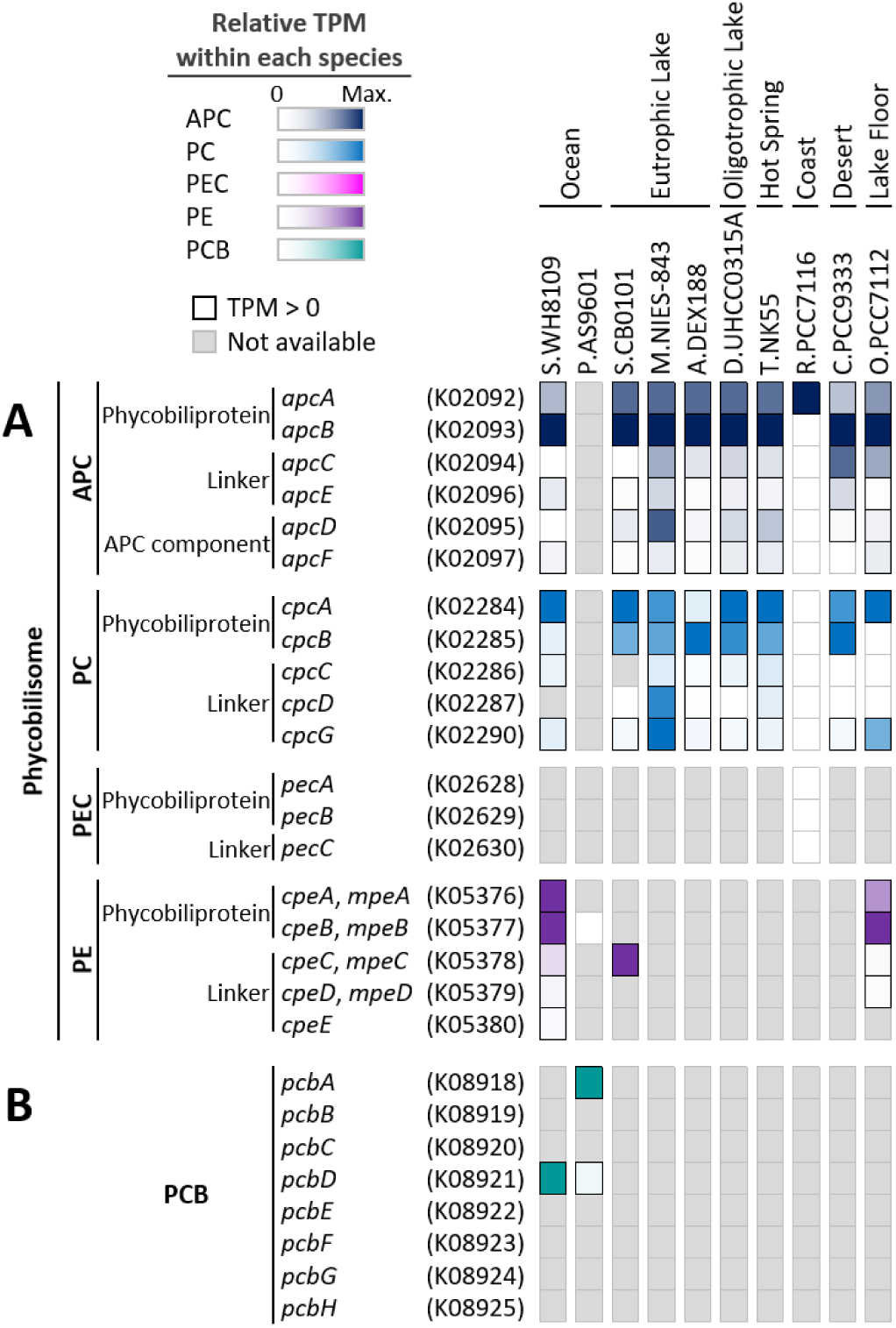
Heatmaps showing the expression profiles of genes encoding the light-harvesting antenna proteins in the daytime samples of the putative dominant species. (A) Genes encoding phycobiliproteins including allophycocyanin (APC), phycocyanin (PC), phycoerythrocyanin (PEC), and phycoerythrin (PE), as well as phycobilisome linker proteins; and (B) genes encoding divinyl chlorophyll *a*/*b*-binding (Pcb) proteins. When multiple genes were assigned the same KO identifier, their mean TPM was used.

In general, phycobiliprotein subunits assemble into the phycobilisome architecture via linkages mediated by their cognate linker proteins (Liu et al. 2005; Stadnichuk et al. 2015). In S.CB0101, however, expression of *cpe*/*mpeC*, which is annotated as a PE-associated rod linker, was detected even though its phycobilisome rods are composed solely of PC (Fig. 4A). Because S.CB0101 lacks *cpcC*, which encodes the PC-associated rod linker, Cpe/MpeC might function as an alternative linker for PC assembly. It should be noted, however, that S.CB0101 was designated as a putative dominant species based on mRNA read mapping rates and may not represent the actual dominant species in the environment. Thus, the absence of *cpcC* and *cpe*/*mpeBA*, which encode PE, together with the presence of *cpe*/*mpeC*, in the genome of S.CB0101 might not reflect the genomic composition of the actual dominant species in that environment.

Most strains of the genus *Prochlorococcus*, including P.AS9601, use divinyl chlorophyll *a*/*b*-binding (Pcb) proteins as their light-harvesting antennae (Hess et al. 2001). At least eight distinct *pcb* genes (*pcbA* through *pcbH*) have been identified in *Prochlorococcus*, and the number of these genes varies from one to eight among species (Ting et al. 2009). P.AS9601 possesses *pcbA* and *pcbD*, of which *pcbA* was expressed at a high level in daytime samples, being included among the highly expressed genes (Fig. 4B and Table S5). In contrast, the TPM of *pcbD* in daytime samples was approximately one-tenth that of *pcbA*. This distinct expression pattern of *pcb* genes in P.AS9601 likely reflected the functional differences between these antenna proteins. In *Prochlorococcus marinus* str. MIT 9313, *pcbA* is constitutively expressed, and PcbA functions as a light-harvesting antenna associated with the PSII dimer, with four PcbA subunits on each side. On the other hand, expression of *pcbD* is induced under iron-stress conditions, and PcbD forms a large ring of 18 subunits surrounding the PSI trimer (Bibby et al. 2003; Ting et al. 2009). Although *Prochlorococcus* lacks phycobilisomes, some strains harbor *cpeB*, which encodes the β subunit of PE, and expression of this gene has been reported in some cases (Hess et al. 1996; Steglich et al. 2005). In this study, expression of *cpeB* was not detected in P.AS9601, despite its presence in the genome (Fig. 4A).

### 5. Diurnal expression patterns of genes encoding PSII and PSI reaction center subunits

We investigated the diurnal expression patterns of the PSII and PSI reaction center genes, *psbA* and *psaA*, respectively, both of which were highly expressed in all putative dominant species (Figs. 3A and 3B). Across the putative dominant species, *psbA* expression increased during daylight and peaked around noon, whereas *psaA* expression increased later and peaked around sunset (Fig. 5A). This phase shift in diurnal expression pattern of *psbA* and *psaA* is consistent with previous findings from a metatranscriptomic analysis of a *Microcystis* bloom in a freshwater lake (Davenport et al. 2019). The increase in *psbA* transcription during daytime might reflect the need to replace photodamaged D1 protein during PSII repair. In contrast, PSI is generally less susceptible to photodamage (Lima-Melo et al. 2021), rendering upregulation of *psaA* transcription during daylight unnecessary. As sunset approaches and solar irradiance declines, light-harvesting capacity for PSII may be enhanced through upregulated expression of phycobilisome-related genes (Fig. S4), whereas PSI binding a large proportion of cellular chlorophyll *a* (Pohland et al. 2024) may increase its own abundance through upregulation of its subunit genes to enhance light capture under dim-light conditions.

**Figure 5.**
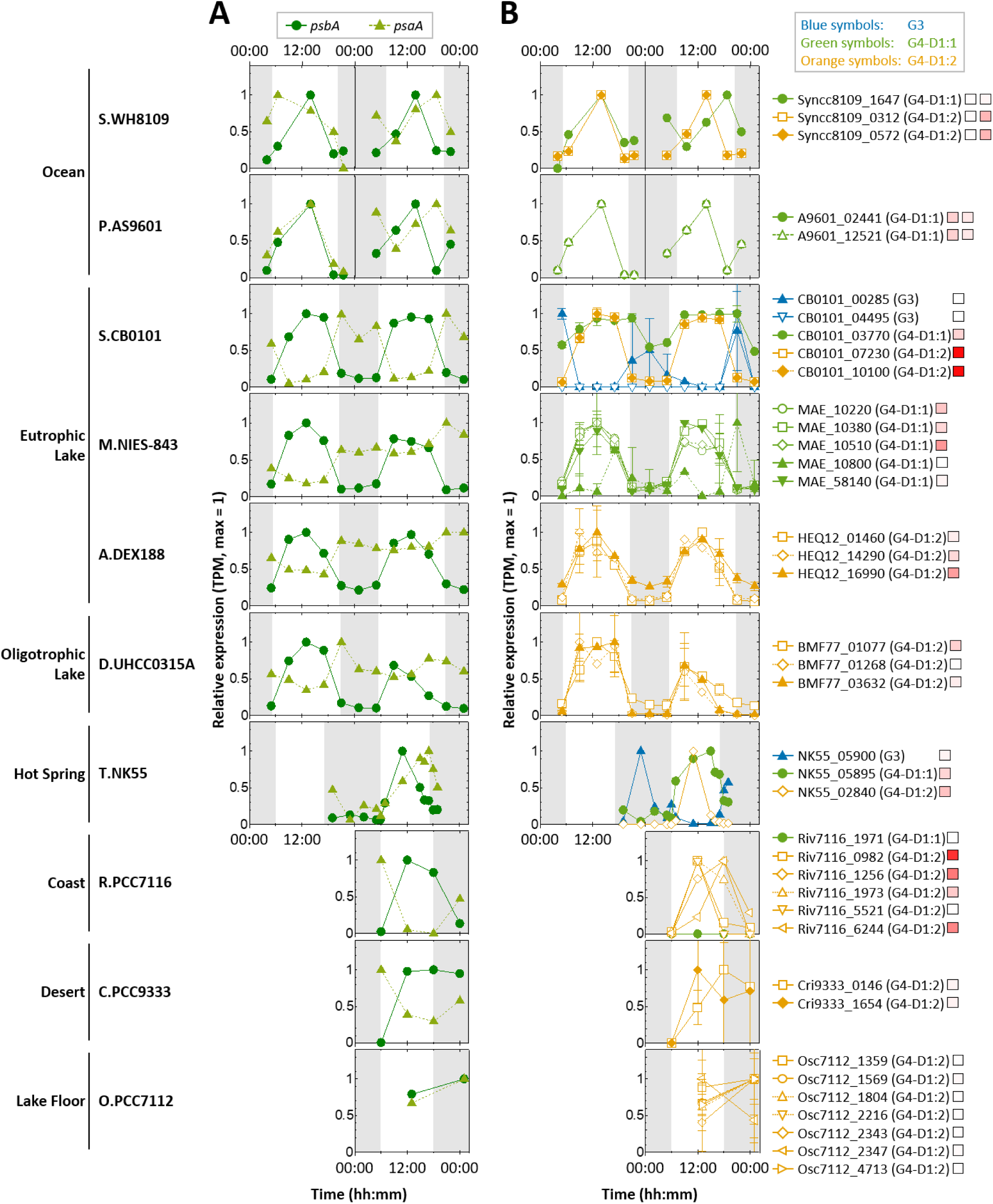
Diurnal expression patterns of (A) *psbA* and *psaA*, assigned KO identifiers K02703 and K02689, respectively, and (B) individual *psbA* copies in each putative dominant species. In each plot, white and gray backgrounds indicate periods with and without sunlight, respectively. TPM values at each sampling time are expressed relative to the maximum TPM across all time-point samples for the corresponding gene, with the maximum set to 1. (A) Circles, *psbA*; triangles, *psaA*. For *psbA* genes, the mean TPM across all assigned KO identifier K02703 in each putative dominant species was calculated at each sampling time, and these mean values were then normalized to the maximum across all time-point samples, which was set to 1. (B) Green symbols, G4-D1:1; orange symbols, G4-D1:2; and blue symbols, G3-D1. The colored squares to the right of each gene ID indicate the maximum TPM for that gene, represented by a continuous color gradient from white (TPM = 0) to red (TPM = 300,000).

The common diurnal expression patterns of *psbA* and *psaA* observed across the putative dominant species suggest regulation by a transcriptional control system universally conserved in cyanobacteria. One possible mechanism is the well-characterized circadian regulatory system in cyanobacteria consisting of the core oscillator formed by KaiA, KaiB, and KaiC and the SasA-RpaA two-component output pathway (Shultzaberger et al. 2015; Hu et al. 2026). However, photosystem reaction center genes were not identified as members of the RpaA regulon (Markson et al. 2013). A more plausible candidate regulator is RpaB, a master transcriptional regulator of photosynthesis-related genes (Wilde and Hihara 2016; Riediger et al. 2018). In model species such as *Synechocystis* sp. PCC 6803, RpaB binds to the high-light regulatory 1 (HLR1) motif under low-light conditions. Under these conditions, RpaB acts as a repressor of *psbA*, whose HLR1 motif lies near the transcription start site (TSS; −38 to +23 nt relative to the TSS), whereas it acts as an activator of *psaA*, whose HLR1 motif lies upstream of the core promoter (−66 to −45 nt relative to the TSS) (Riediger et al. 2018). Such position-dependent transcription regulation by RpaB relative to the promoter architecture may drive the phase shift in diurnal expression patterns of *psbA* and *psaA* observed across the putative dominant species (Fig. 5A). RpaB has been reported to be present in all cyanobacteria with fully sequenced genomes (Riediger et al. 2018). We found the HLR1 motifs within 400 bp upstream of the start codons of both *psbA* and *psaA* in all ten putative dominant species designated in this study (Fig. S5). The HLR1 motifs in the *psbA* upstream regions were concentrated around 100 bp upstream of the start codons (Fig. S5A), whereas those in the *psaA* upstream regions tended to occur farther upstream (Fig. S5B), which is consistent with the observation by Riediger et al. (2018). HLR1 motifs have also been found upstream of the core promoter of the *cpcBA* operon encoding PC of phycobilisomes (Riediger et al. 2018). Consistent with this, *cpcBA* expression, like that of *psaA*, peaked around sunset across the putative dominant species (Fig. S4). Taken together, these observations suggest that, irrespective of environmental conditions or cyanobacterial taxa, RpaB-mediated transcriptional regulation may contribute to the diurnal expression patterns of major, highly expressed photosynthesis-related genes, including genes encoding photosystems and phycobilisome subunits.

Since the existence of five D1 protein isoforms (G0-G4) has been reported in cyanobacteria (Cardona et al. 2015), we evaluated the diurnal expression patterns of the *psbA* genes encoding each isoform individually (Fig. 5B). Across the putative dominant species, expression of the *psbA* genes encoding G4-type D1 (G4-D1), the major D1 isoform involved in oxygen evolution and conserved also in land plants (Cardona et al. 2015), was consistently induced by daylight (Fig. 5B). G4-D1 comprises two variants: D1:1 and high-light tolerant D1:2, in which glutamine at position 130 is substituted with glutamate (Golden et al. 1986; Clarke et al. 1993). In S.WH8109, S.CB0101, T.NK55, and R.PCC7116, all of which possess both variants, the maximum TPM of the gene encoding D1:2 was higher than that of the gene encoding D1:1 (Fig. 5B). In T.NK55, expression of the D1:1 gene was induced soon after sunrise, whereas the D1:2 gene was barely induced at dawn and instead exhibited a pronounced expression peak around noon (11:00). This transcriptional activation of the D1:2 gene around midday may mitigate PSII photodamage. In addition to the G4-type, S.CB0101 and T.NK55 possess the G3-type D1 isoform (G3-D1), whose encoding gene is induced under microoxic conditions (Summerfield et al. 2008; Sicora et al. 2009). In both species, expression of the G3-D1 gene was induced during the dark period (Fig. 5B), likely reflecting the formation of a hypoxic environment caused by the cessation of photosynthetic oxygen evolution and ongoing oxygen consumption via respiration at night. Why G3-D1 was required and induced during the night remains an open question, and its physiological significance warrants further investigation.

### 6. Functional characteristics of *psbA* co-expressed genes

To elucidate the shared and unique features of gene sets induced by increased light intensity across diverse cyanobacteria inhabiting distinct natural environments, we cataloged genes co-expressed with *psbA* encoding G4-D1, whose expression consistently increased during daylight across the putative dominant species (Fig. 5B). For this analysis, we focused on M.NIES-843, A.DEX188, D.UHCC0315A, and T.NK55, in which the proportion of detected genes exceeded 50% of the total genome-encoded genes (Fig. S2). In T.NK55, because the two *psbA* genes encoding D1:1 and D1:2 showed different patterns of daylight-associated induction (Fig. 5B), we analyzed co-expression separately using each gene as a reference. Nearly half of the co-expressed genes identified by this approach lacked an assigned KO identifier (Table S6; 250 of 550 gene instances in total). We therefore classified all genes from the putative dominant species into orthogroups and searched for orthogroups shared among the *psbA*-co-expressed gene sets, including those of unknown function. As a result, nine *psbA*-co-expressed orthogroups were identified that were shared by at least three of the four putative dominant species (Table 3).

**Table 3.** Shared orthogroups of psbA-co-expressed genes across putative dominant species.

| **Orthogroup** | ***Synechocystis* sp. PCC 6803 gene ID** | **KO identifier** | **KO description** | **M.NIES-843** | **A.DEX188** | **D.UHCC0315A** | **T.NK55 (D1:1)** | **T.NK55 (D1:2)** |
|---|---|---|---|---|---|---|---|---|
| OG0000038 | slr1181 (*psbA1*),<br>slr1311 (*psbA2*),<br>sll1867 (*psbA3*) | K02703 | psbA; photosystem II P680 reaction center D1 protein [EC:1.10.3.9] | 3 (5) | 2 (3) | 2 (3) | 1 (3) | 1 (3) |
| OG0000200 | sll0849 (*psbD1*),<br>slr0927 (*psbD2*) | K02706 | psbD; photosystem II P680 reaction center D2 protein [EC:1.10.3.9] | 2 (2) | 1 (2) | 2 (2) | 2 (2) | 2 (2) |
| OG0000652 | sll0851 (*psbC*) | K02705 | psbC; photosystem II CP43 chlorophyll apoprotein | 1 (1) | 0 (1) | 1 (1) | 1 (1) | 1 (1) |
| OG0000026 | ssl2542 (*hliA*),<br>ssr2595 (*hliB*),<br>ssl1633 (*hliC*),<br>ssr1789 (*hliD*) | N/A | N/A | 1 (3) | 0 (2) | 1 (3) | 2 (4) | 1 (4) |
| OG0000070 | sll0306 (*sigB*),<br>sll0184 (*sigC*),<br>sll2012 (*sigD*) | K03087 | rpoS; RNA polymerase nonessential primary-like sigma factor | 1 (3) | 0 (3) | 1 (3) | 1 (3) | 1 (3) |
| OG0000151 | slr1963 (*ocp*) | N/A | N/A | 0 (1) | 1 (1) | 1 (1) | 0 (2) | 1 (2) |
| OG0000120 | slr0331 (*ndhD1*),<br>slr1291 (*ndhD2*) | K05575 | ndhD; NAD(P)H-quinone oxidoreductase subunit 4 [EC:7.1.1.2] | 1 (2) | 1 (3) | 1 (3) | 1 (2) | 1 (2) |
| OG0000547 | sll0185 | K09942 | K09942; uncharacterized protein | 1 (1) | 1 (1) | 1 (1) | 1 (1) | 1 (1) |
| OG0000257 | slr1053 | N/A | N/A | 1 (2) | 1 (1) | 1 (1) | 0 (0) | 0 (0) |

Values represent the number of *psbA* co-expressed genes in the orthogroup (total number of genes in the orthogroup). An orthogroup was considered common when it contained at least one *psbA* co-expressed gene in at least three of the four putative dominant species. For T.NK55, the orthogroup was counted as present if it was identified in either the D1:1- or D1:2-based co-expression analysis.

Three of the nine orthogroups shared across the *psbA*-co-expressed gene sets (Table 3) corresponded to genes encoding PSII subunits, including *psbA*, *psbD*, and *psbC*. In addition, the following three orthogroups were listed: OG0000026, corresponding to genes encoding high-light-inducible proteins (HLIPs), which are chlorophyll-binding proteins that protect PSII assembly intermediates under high-light conditions (Iwai et al. 2024); OG0000070, corresponding to genes encoding the group 2 sigma factors SigB, SigC and SigD (Srivastava et al. 2020); and OG0000151, corresponding to genes encoding the orange carotenoid protein (OCP), which dissipates excess light energy absorbed by phycobilisomes as heat, thereby decreasing the transfer of excitation energy to photosystem reaction centers (Kerfeld and Sutter 2024). Phylogenetic analysis confirmed that the group 2 sigma factor gene found to be co-expressed with *psbA* in the three putative dominant species other than A.DEX188 encoded SigD (Fig. S6), whose expression is induced by high light and which is essential for high-light acclimation (Imamura et al. 2003; Pollari et al. 2008). These results suggest that genes encoding HLIPs, SigD, and OCP, which have been shown to play important roles in high-light acclimation in laboratory-grown model species, commonly exhibit daylight-associated induction together with the G4-D1 gene across cyanobacteria inhabiting different natural environments.

Another orthogroup shared across *psbA*-co-expressed gene sets of the putative dominant species was OG0000120, corresponding to *ndhD* genes encoding subunits of the NDH-1 complex (Table 3). In *Synechocystis* sp. PCC 6803, different sets of NdhD and NdhF isoforms are known to be incorporated into several types of NDH-1 complexes with distinct functions (Ma et al. 2021). OG0000120 contained *ndhD1* and *ndhD2*, which encode subunits specific to the NDH-1L and NDH-1L’ complexes, respectively, both of which participate in respiratory and PSI cyclic electron transport. In M.NIES-843, A.DEX188, and D.UHCC0315A, *ndhD2* was identified as a *psbA*-co-expressed gene (Table S6), whereas *ndhD1* tended to be induced during dark periods (Fig. S7). These distinct expression patterns of *ndhD1* and *ndhD2* suggest that the NDH-1L and NDH-1L’ complexes are not physiologically redundant, although the exact functional differences between these complexes remain unclear. In T.NK55, on the other hand, *ndhD1* expression also increased during daylight (Fig. S7), and both *ndhD2* (NK55_05770) and *ndhD1* (NK55_10110) were identified as *psbA*-co-expressed genes (Tables S6 and S7). Given that a transition from 30°C to 55°C stimulates PSI cyclic electron transport in *Synechocystis* sp. PCC 6803 (Ma et al. 2008), the daylight-dependent induction of *ndhD1* observed specifically in T.NK55 of the Hot Spring dataset might reflect an elevated cellular demand for NDH-1L-mediated PSI cyclic electron transport under high-temperature environments.

Two of the nine orthogroups shared across *psbA*-co-expressed gene sets of the putative dominant species were of unknown function: OG0000547 and OG0000257 (Table 3). Genes assigned to OG0000547 were present in all 159 cyanobacterial strains analyzed in this study (Fig. S8). In *Synechocystis* sp. PCC 6803, the OG0000547 gene *sll0185* encodes a protein containing a Rho_N domain (Rho termination factor, N-terminal domain), and expression of *sll0185* has been reported to be induced under hexane and osmotic stress conditions but repressed under nitrogen-starvation conditions (Qiao et al. 2013). Genes assigned to OG0000257 were present in 118 of the 159 strains (74%) but were absent from certain lineages, including *Thermosynechococcus*, *Synechococcus elongatus*, marine *Synechococcus*, and *Prochlorococcus* (Fig. S8). These two uncharacterized orthogroups are particularly noteworthy as they may participate in yet-undiscovered high-light acclimation mechanisms that are widely conserved among cyanobacteria.

Species-specific *psbA*-co-expressed genes were also identified in A.DEX188 and D.UHCC0315A. In A.DEX188, the number of *psbA*-co-expressed genes was relatively small (14 genes) compared to the other three putative dominant species, but four of them encoded gas vesicle structural proteins, Gvps (Table S6). Among the other three species, only D.UHCC0315A had a single *gvp* gene among its *psbA*-co-expressed genes. Thus, the high frequency of *gvp* genes in the *psbA*-co-expressed gene set was a feature unique to A.DEX188. In contrast, D.UHCC0315A showed a high frequency of chaperone-related genes, and the KEGG PATHWAY category "Folding, sorting and degradation" was significantly enriched in the *psbA*-co-expressed genes only in this species (Fig. S9). Furthermore, multiple genes involved in protein quality control of the photosynthetic machinery were identified among the genes co-expressed with *psbA*, including those encoding the FtsH protease, which is involved in the degradation of photodamaged D1 (Krynická and Komenda 2024), and the Clp protease, which participates in phycobilisome degradation (Bouchnak and van Wijk 2021) (Table S6).

D1:2-encoding *psbA* of T.NK55 showed a pronounced expression peak at 11:00 (Fig. 5B). Namely, the TPM ratio at 11:00 to that at 7:00 (TPM_11:00_/TPM_7:00_) was approximately 20 for D1:2-encoding *psbA*, compared with approximately 1.5 for D1:1-encoding *psbA* (Table S7). The D1:2-encoding *psbA* co-expressed genes in T.NK55 were also characterized by high TPM_11:00_/TPM_7:00_ ratios (Table S7). Notably, four of the ten D1:2-encoding *psbA* co-expressed genes with the highest TPM_11:00_/TPM_7:00_ ratios were *ndh* genes: *ndhD2* mentioned above; *ndhF3* (NK55_11095) and *ndhD3* (NK55_11100), which encode subunits specific to the NDH-1MS complex involved in high-affinity CO_2_ uptake; and *ndhK* (NK55_10035), which encodes a subunit common to all NDH-1 complexes (Table S7). These results suggest that, in the Hot Spring dataset, the demand might be elevated not only for NDH-1L-mediated PSI cyclic electron transport but also for high-affinity CO_2_ uptake mediated by NDH-1MS. In T.NK55, genes encoding common NDH-1 subunits other than *ndhK* also exhibited pronounced increases in expression at 11:00 (Fig. S10), and this induction of the *ndh* genes may be reflected in the significant enrichment of the KEGG PATHWAY category "Oxidative phosphorylation" among highly expressed genes specifically at 11:00 (Fig. S3). A similar set of *ndh* genes also showed pronounced induction in a genome-wide expression analysis of *Synechocystis* sp. PCC 6803 following a shift from low to high light intensity (Hihara et al. 2001).

### 7. Expression profiles of genes involved in transcriptional regulation

To characterize features of transcriptional regulatory mechanisms involved in environmental responses across different environments and cyanobacterial species, we examined the expression profiles of genes assigned to the "Two-component system" and "Transcription factors" categories in the KEGG BRITE database (Fig. 6). Because these categories included many genes whose TPM values were zero at some time points, we calculated the mean TPM across all time points for each putative dominant species to compare their overall expression profiles. In T.NK55, in which the proportion of detected genes relative to the total number of genes was markedly higher than in the other putative dominant species (> 90%; Figs. 2 and S2), more than half of the genes in each category had mean TPM values greater than 100 (Fig. S11). In all other putative dominant species, more than half of the genes in these categories had mean TPM values below 100 (Fig. S11), suggesting generally low basal expression levels of the genes assigned to these categories.

**Figure 6.**
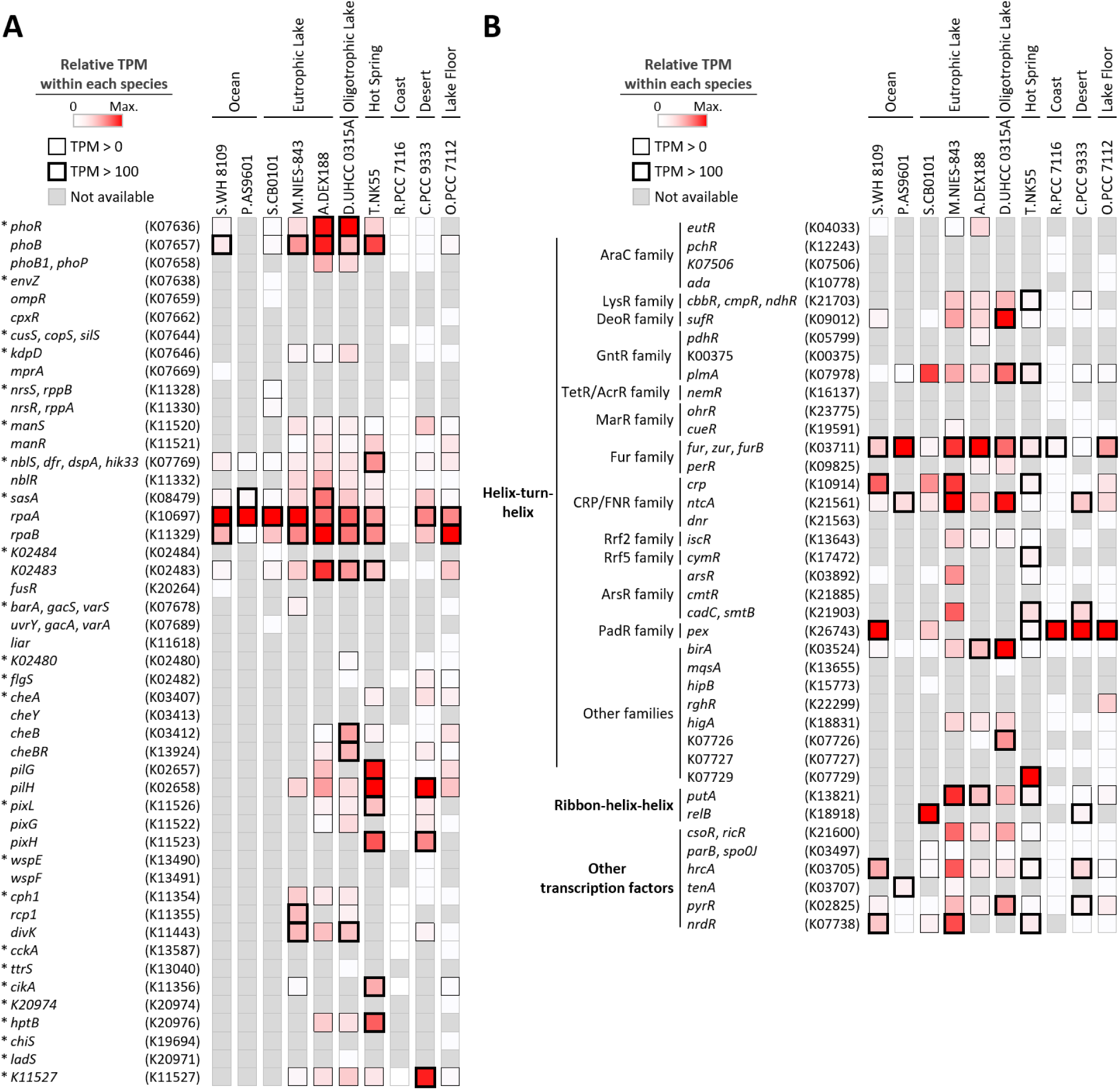
Heatmaps showing the expression profiles of genes assigned to "Two-component system" (A) and "Transcription factors" (B) of the KEGG BRITE database. In panel A, asterisks indicate genes annotated as histidine kinases. The mean TPM across samples collected at all time points is shown for each putative dominant species. When multiple genes were assigned the same KO identifier, their mean TPM was used.

Among the genes encoding sensor histidine kinases and response regulators classified in "Two-component system", either *rpaA* or *rpaB*, or both had mean TPM values exceeding 100 in all putative dominant species except R.PCC7116, in which all genes in this category had TPM values of zero (Fig. 6A). In particular, *rpaA* had the highest mean TPM among the genes in this category in S.WH8109, P.AS9601, S.CB0101, and M.NIES-843, whereas *rpaB* had the highest mean TPM in A.DEX188 and O.PCC7112.

Some genes in "Two-component system" showed species-specific high expression, with TPM values exceeding those of *rpaA* and *rpaB*. In T.NK55 of the Hot Spring dataset and C.PCC9333 of the Desert dataset, *pilH* had the highest TPM among genes in this category (Fig. 6A). In *Synechocystis* sp. PCC 6803, PilH (Slr1042) has been reported to be a response regulator involved in type IV pilus biogenesis and motility (Yoshihara et al. 2002). The relatively high expression of *pilH* in T.NK55 and C.PCC9333, representing species inhabiting surface-associated mats or crust habitats, might reflect the importance of motility on solid surfaces in their habitats. In D.UHCC0315A of the Oligotrophic Lake dataset, *phoR* showed the highest expression among genes classified in "Two-component system" (Fig. 6A), and relatively high expression of *phoR* and/or *phoB* was also observed in A.DEX188 of the Eutrophic Lake dataset and T.NK55 of the Hot Spring dataset (Fig. 6A). In *Synechocystis* sp. PCC 6803, a two-component signal transduction system comprising SphS (Sll0337), a homolog of the histidine kinase PhoR, and SphR (Slr0081), a homolog of the response regulator PhoB, regulates the expression of genes encoding proteins such as alkaline phosphatase in response to inorganic phosphate deficiency (Hirani et al. 2001; Suzuki et al. 2004). These results suggest that the phosphate-responsive two-component system might be important in diverse environments.

In "Transcription factors", the gene encoding Fur, which regulates iron uptake and related processes in an iron concentration-dependent manner (Andrews et al. 2003), exhibited TPM values greater than 100 in eight of the ten putative dominant species (Fig. 6B). In particular, *fur* showed the highest TPM among genes in this category in P.AS9601 and A.DEX188. In addition, a gene encoding Pex, which binds to the promoter region of *kaiA* and acts as its transcriptional repressor in *Synechococcus elongatus* PCC 7942 (Kutsuna et al. 1998; Arita et al. 2007; Mackey and Golden 2007), was conserved in six of the ten putative dominant species, with five of them exhibiting a TPM value greater than 100 (Fig. 6B). In S.WH8109, R.PCC7116, C.PCC9333, and O.PCC7112, *pex* showed the highest TPM among genes in "Transcription factors". These results suggest that *fur* and *pex* were commonly expressed at relatively high levels across putative dominant species from different environments.

In M.NIES-843, D.UHCC0315A, and T.NK55, different genes in "Transcription factors" showed relatively high expression, with TPM values exceeding those of *fur* or *pex* (Fig. 6B). *ntcA*, encoding the global nitrogen regulator (Forchhammer and Selim 2020) in M.NIES-843, and *birA*, encoding the repressor of biotin biosynthesis (Mills et al. 2020) in D.UHCC0315A showed TPM values comparable to those of *fur* regardless of the time period (Fig. S12). In T.NK55, TPM for the gene assigned the KO identifier K07729 which is annotated as a "putative transcriptional regulator" increased during dark periods, reaching values markedly higher than those of other transcription factor genes at night (Fig. S13).

Taken together, even in putative dominant species with a low proportion of detected genes relative to total genome-encoded genes (<20%), expression of specific transcriptional regulatory genes could be reliably detected, enabling us to identify regulatory genes commonly or specifically expressed across different environments and cyanobacterial species. Our analysis demonstrated that metatranscriptomic datasets can be used to examine the expression profiles of not only highly expressed core photosynthetic genes, but also low-abundance genes encoding transcriptional regulatory machinery.

## Conclusion

In this study, by mapping metatranscriptomic reads to the genome sequences of phylogenetically diverse cyanobacteria, we systematically compared the gene expression profiles across multiple natural environments and taxa using a unified analytical framework. These analyses revealed responses shared among phylogenetically diverse cyanobacteria in distinct natural environments, such as the phase shift in diurnal expression patterns of PSII and PSI reaction center genes possibly regulated by RpaB (Figs. 5 and S5), and high-light acclimation involving OCP, HLIPs, and SigD (Table 3). Conversely, we also captured environment- or species-specific expression profiles, such as daylight-dependent induction of *ndhD1* of NDH-1L in the hot spring cyanobacterium and of *gvp* encoding gas vesicle structural proteins in the eutrophic lake cyanobacterium (Fig. S6). Some putative dominant species also showed expression profiles that have not been documented in laboratory studies, including possibly higher expression levels of small subunit genes than reaction center genes of photosystems (Fig. 3A, B) and expression of a gene encoding a PE-associated rod linker protein in a putative dominant species lacking PE (Fig. 4A). These distinct transcriptional responses may support physiological acclimation or adaptation to the complexities of natural habitats. Notably, along with known high-light acclimation genes, uncharacterized genes showing daylight-dependent induction were identified through co-expression analysis with *psbA* (Table 3), pinpointing candidate factors that may participate in novel, yet-uncharacterized high-light acclimation mechanisms. Such candidate genes identified in natural populations can in turn be functionally characterized in model strains, and such a cycle between field observations and laboratory experiments will deepen our understanding of how cyanobacteria thrive in fluctuating natural environments. Finally, even in putative dominant species with a low proportion of detected genes, the expression of low-abundance regulatory genes, such as those encoding two-component systems and transcription factors, could be detected, revealing regulators commonly expressed across species, such as *rpaA*, *rpaB*, *fur*, and *pex*, as well as species-specific ones (Fig. 6). The shared and environment-or species-specific features of expression patterns of photosynthesis-related genes and regulatory genes observed in this study provide valuable insights into the fundamental mechanisms governing environmental acclimation and adaptation in cyanobacteria. As metatranscriptomic data continue to accumulate in public databases, the approach presented here can be applied to a wider range of environments and taxa.

## Materials and Methods

### Acquisition and quality control of metatranscriptomic data

Data shown in Table S1 were obtained from the SRA of NCBI (Leinonen et al. 2011). The obtained data were converted to FASTQ format using SRA Toolkit 2.8.0. The resulting FASTQ files were subjected to quality control and adapter sequence removal using fastp 0.23.3 (Chen 2023), followed by removal of rRNA reads using SortMeRNA 2.1b (Kopylova et al. 2012). A SILVA-derived database containing bacterial and archaeal 16S and 23S rRNAs and eukaryotic 18S and 28S rRNAs was used as the reference for rRNA removal. Reads unmapped to the rRNA reference sequences were treated as mRNA reads and used for subsequent analyses.

### Determination of putative dominant cyanobacterial species and quantification of transcript levels

To identify putative dominant cyanobacterial species, 169 of the 174 strains with genome information registered in the KEGG database (Kanehisa et al. 2025; https://www.genome.jp/kegg/) as of August 1, 2024 were selected as candidates after excluding the five substrains of *Synechocystis* sp. PCC 6803. Among these, 164 strains harboring all four genes assigned KO identifiers for PSII and PSI reaction center genes (*psbA*, K02703; *psbD*, K02706; *psaA*, K02689; *psaB*, K02690) were selected. Whole-genome sequence files (FASTA) and gene annotation files (GFF3) of 160 strains could be obtained from either GenBank or RefSeq. Among 160 strains, *Nodularia spumigena* UHCC 0039 was excluded because the number of genes reported in GenBank was substantially lower than that in RefSeq, corresponding to approximately 51% of the RefSeq gene count, suggesting that the GenBank gene annotation may be incomplete. The remaining 159 strains (Table S2) were used for subsequent analyses.

For each strain, an RSEM 1.3.3 (Li and Dewey 2011) reference was constructed using the whole-genome sequence file and the GFF3-format gene annotation file. The mRNA reads from each sample were independently mapped to the RSEM reference of each of the 159 strains using Bowtie2 2.5.4 (Langmead and Salzberg 2012) with a seed length of 25 bp and an allowed mismatch rate of 0.1. For the samples marked with an asterisk in Table S1, the mapping rate was calculated as the proportion of mRNA reads mapped to each reference relative to the total number of mRNA reads in each sample. For each environment, the strain showing the highest mapping rate was selected as the putative dominant species. When strains with mapping rates greater than 0% were distributed across multiple clades in the cyanobacterial species tree (Fig. 1), the strain with the highest mapping rate was selected from each clade.

The mRNA reads from the remaining samples without asterisks in Table S1 were also mapped to the RSEM reference of selected putative dominant species, and transcript levels were quantified for each sample. Transcript levels were calculated as Transcripts Per Million (TPM).

### Orthogroup inference and construction of cyanobacterial phylogenetic tree

Orthogroups were inferred from all protein sequences of the 159 cyanobacterial strains using OrthoFinder 2.2.7 (Emms and Kelly 2015). From the 180 identified single-copy orthologs, twelve for which the longest sequence was more than twice as long as the shortest sequence were excluded because their amino acid sequence lengths varied substantially among strains. To prepare a concatenated alignment for phylogenetic analysis, the amino acid sequences of the remaining 168 single-copy orthologs across the 159 strains were aligned using MAFFT v7.525 (Katoh and Standley 2013), and poorly aligned regions were trimmed using trimAl v1.5.rev1 (Capella-Gutiérrez et al. 2009). The 126 orthologs whose post-trimming alignment lengths exceeded 100 amino acids and also exceeded 80% of their respective pre-trimming alignment lengths were selected for phylogenetic tree construction. The 126 trimmed alignments were concatenated, and a maximum-likelihood phylogenetic tree of cyanobacteria was inferred using IQ-TREE 2.4.0 (Minh et al. 2020), with each ortholog initially defined as a separate partition. The MFP+MERGE option was used for model selection and partition merging, and branch support was assessed using 1,000 ultrafast bootstrap replicates. *Gloeobacter violaceus* PCC 7421, *Gloeobacter kilaueensis* JS1, and *Gloeobacter morelensis* MG652769 were used as outgroups. The phylogenetic tree was visualized using iTOL v7 (Letunic and Bork 2024).

### KEGG PATHWAY enrichment analysis

For each time-point sample of each putative dominant species, genes with TPM values greater than 0 (detected genes) were ranked in descending order of TPM. The top 10% of detected genes were selected as highly expressed genes, with the number of selected genes rounded to the nearest integer when necessary. Gene functional classifications were assigned based on KO identifiers, using the second-level categories in the KEGG PATHWAY hierarchy. For the "Energy metabolism" category, the third-level subcategories were also analyzed. All genes in the genome of each putative dominant species were defined as the gene universe (background gene set), including genes without assigned KO identifiers. Fold enrichment for each KEGG PATHWAY category was calculated as follows: Fold enrichment = (*k*/*n*)/(*m*/*N*), where *N* is the total number of genes in the reference genome, *m* is the number of genes assigned to a given KEGG PATHWAY category among the *N* genes, *n* is the total number of the highly expressed genes, and *k* is the number of genes assigned to a given KEGG PATHWAY category among the *n* genes. The over-representation of each KEGG PATHWAY category among the highly expressed genes relative to the gene universe was assessed using a one-sided Fisher’s exact test. The resulting P values were adjusted separately for each time-point sample across all KEGG PATHWAY categories tested using the Benjamini-Hochberg method. Categories with a fold enrichment > 1 and a false discovery rate (FDR) < 0.05 were considered significantly enriched.

### In silico identification of putative HLR1 motifs in the upstream regions of *psbA* and *psaA* genes

For each putative dominant species, an HLR1 motif model was constructed using MEME (MEME Suite v5.5.9; Bailey et al. 2015) based on 18 previously reported HLR1 sequences compiled by Riediger et al. (2018), for which RpaB binding had been experimentally confirmed *in vivo* or *in vitro*. MEME was run using the OOPS (One Occurrence Per Sequence) model to identify a single motif with a fixed width of 18 bp. A zero-order Markov background model generated from the whole-genome sequence of each putative dominant species using fasta-get-markov in the MEME Suite was used for motif discovery and subsequent scanning. The resulting HLR1 motif model was used to scan the 400-bp regions upstream of the start codons of *psbA* and *psaA* in each putative dominant species using FIMO (MEME Suite v5.5.9; Grant et al. 2011). Motif matches with P values < 1 × 10^-3^ were identified as putative HLR1 motifs.

### Co-expression analysis with *psbA*

Co-expression analysis with *psbA* was performed for putative dominant species in which the mean proportion of detected genes relative to the total number of genes encoded in the genome exceeded 50% across all time-point samples. The *psbA* gene encoding G4-D1 was used as the reference. When multiple copies of the G4-D1-encoding *psbA* were present in a putative dominant species, the copy with the highest mean TPM across all time-point samples was selected as the reference. For each putative dominant species, gene expression levels were transformed as log_2_(TPM + 1), and Pearson’s correlation coefficients between the expression profiles of individual genes and that of the selected reference *psbA* were calculated. Genes other than the reference *psbA* with Pearson’s correlation coefficients greater than 0.7 were identified as *psbA* co-expressed genes.

### Phylogenetic analysis of sigma factors

The amino acid sequences of the Group 2 sigma factors assigned to orthogroup OG0000070 in the 10 putative dominant species and *Synechocystis* sp. PCC 6803 were aligned using MAFFT v7.525 (Katoh and Standley 2013) with the –auto option. A maximum-likelihood phylogenetic tree was inferred from the resulting multiple sequence alignment using IQ-TREE 2.4.0 (Minh et al. 2020). The best-fit amino acid substitution model was selected using ModelFinder (MFP). Branch support was assessed using 1,000 ultrafast bootstrap replicates.

## Data availability statement

The metatranscriptomic datasets and cyanobacterial reference genome sequences analyzed in this study were retrieved from publicly available databases. The corresponding accession numbers and database information are provided in Tables S1 and S2, respectively.

## Acknowledgements

This work was supported by NIBB Collaborative Research Program (26NIBB338) to YH. The computation was performed using Research Center for Computational Science, Okazaki, Japan (Project: NIBB, 24-IMS-C391, 25-IMS-C237, and 26-IMS-C256). This study is supported by the Japan Society for the Promotion of Science KAKENHI (grant numbers 21H05040, 23H04960, and 26H02384 to JM and 23H04962 to YH).

## Ethics statement

Not applicable.

## Author contributions

**Takako Ogawa:** investigation, methodology, writing - original draft preparation. **Jun Minagawa:** resources, writing - review and editing, funding acquisition. **Yukako Hihara:** conceptualization, writing - original draft preparation, review and editing, funding acquisition.

