## Supporting Information for "Metatranscriptomic reanalysis reveals common and specific photosynthetic acclimation of cyanobacteria across diverse natural environments"

**Contents of Supporting Information**

Supplementary Figures S1–S13 (PDF)

Supplementary Tables S1–S7 (provided as a separate Excel file)

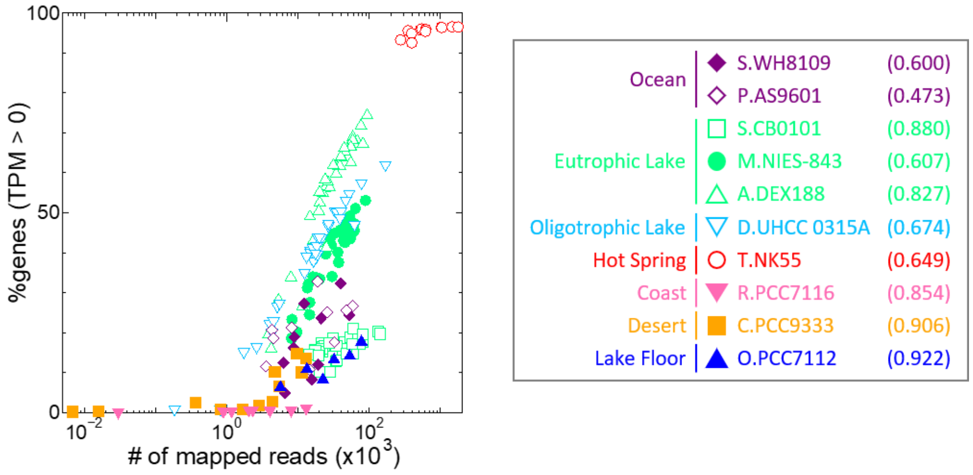


**Figure S1.** Correlation between the number of mRNA reads mapped to the genome of each putative dominant species and the proportion of detected genes (TPM > 0) among all genes encoded in its genome. The value in the parentheses next to the name of each putative dominant species within the box indicates the correlation coefficient for that species.

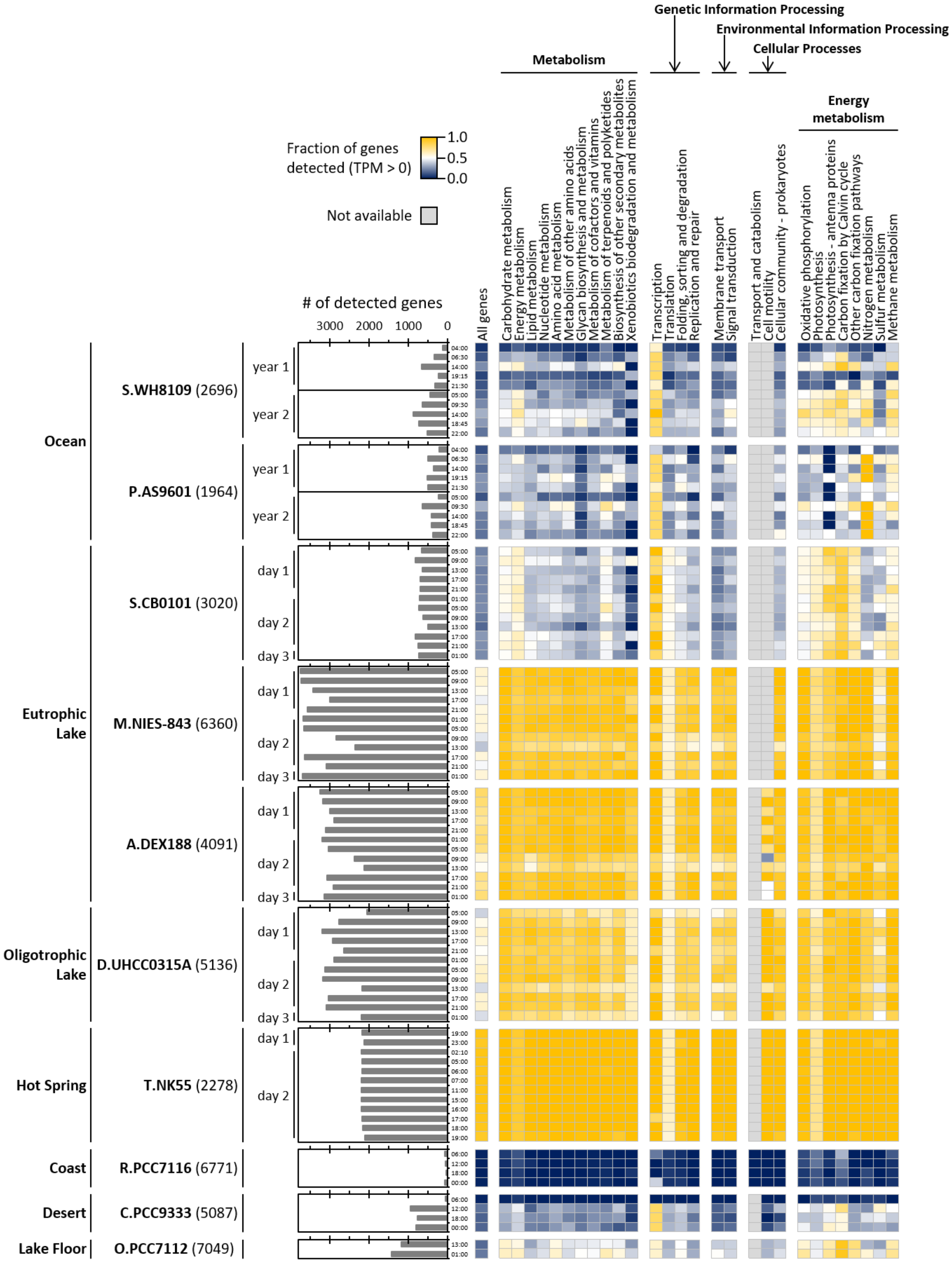


**Figure S2.** Bar chart showing the number of detected genes among all genes encoded in the genome of each putative dominant species, and heatmap showing the proportion of detected genes in each KEGG PATHWAY category. The value in the parentheses to the right of the name of each putative dominant species indicates the total number of genes encoded in its genome.

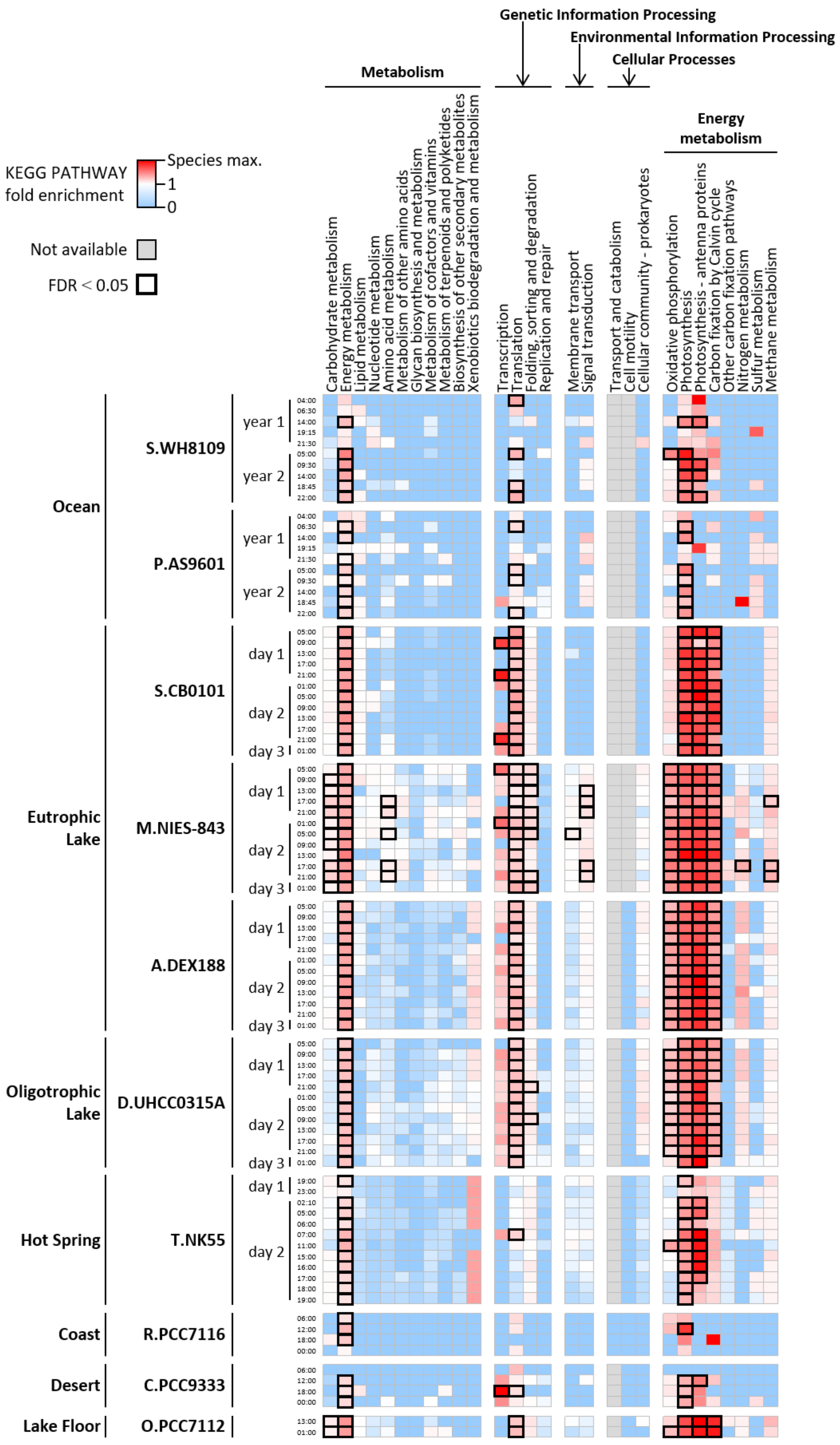


**Figure S3.** Heatmap showing fold enrichment of KEGG PATHWAY categories among highly expressed genes, defined as the top 10% of detected genes ranked by TPM in each sample.

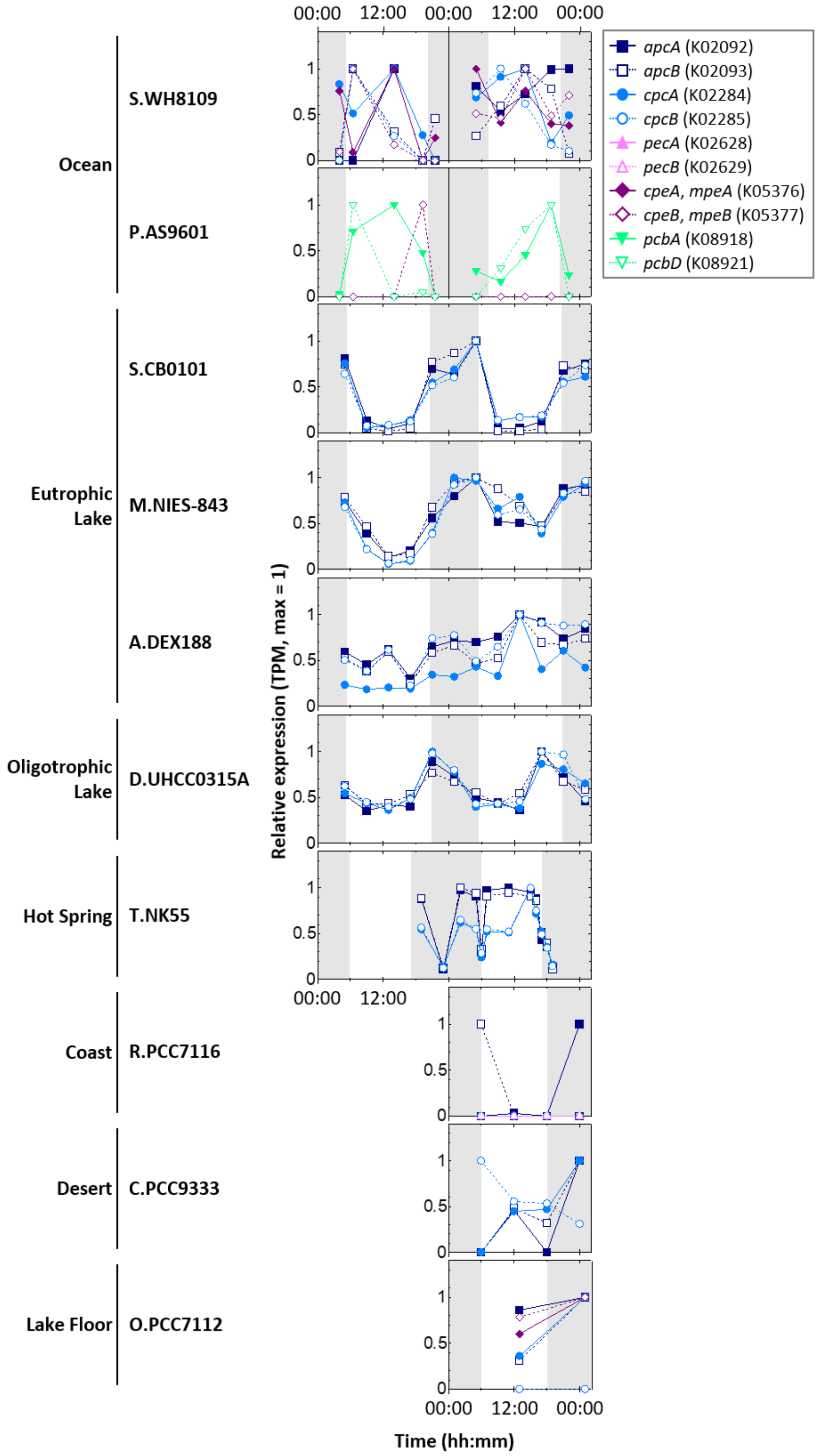


**Figure S4.** Diurnal expression patterns of genes encoding subunits of light-harvesting antenna complexes. In each plot, white and gray backgrounds indicate periods with and without sunlight, respectively. TPM values at each sampling time are expressed relative to the maximum TPM across all time-point samples for the corresponding gene, with the maximum set to 1. When multiple genes were assigned the same KO identifier, their mean TPM was calculated at each sampling time, and the resulting mean values were normalized to the maximum across all time-point samples, which was set to 1. *apcA* (K02092), closed squares; *apcB* (K02093), open squares; *cpcA* (K02284), closed circles; *cpcB* (K02285), open circles; *pecA* (K02628), closed triangles; *pecB* (K02629), open triangles; *cpeA*/*mpeA* (K05376), closed diamonds; *cpeB*/*mpeB* (K05377), open diamonds; *pcbA* (K08918), closed inverted triangles; *pcbD* (K08921), open inverted triangles.

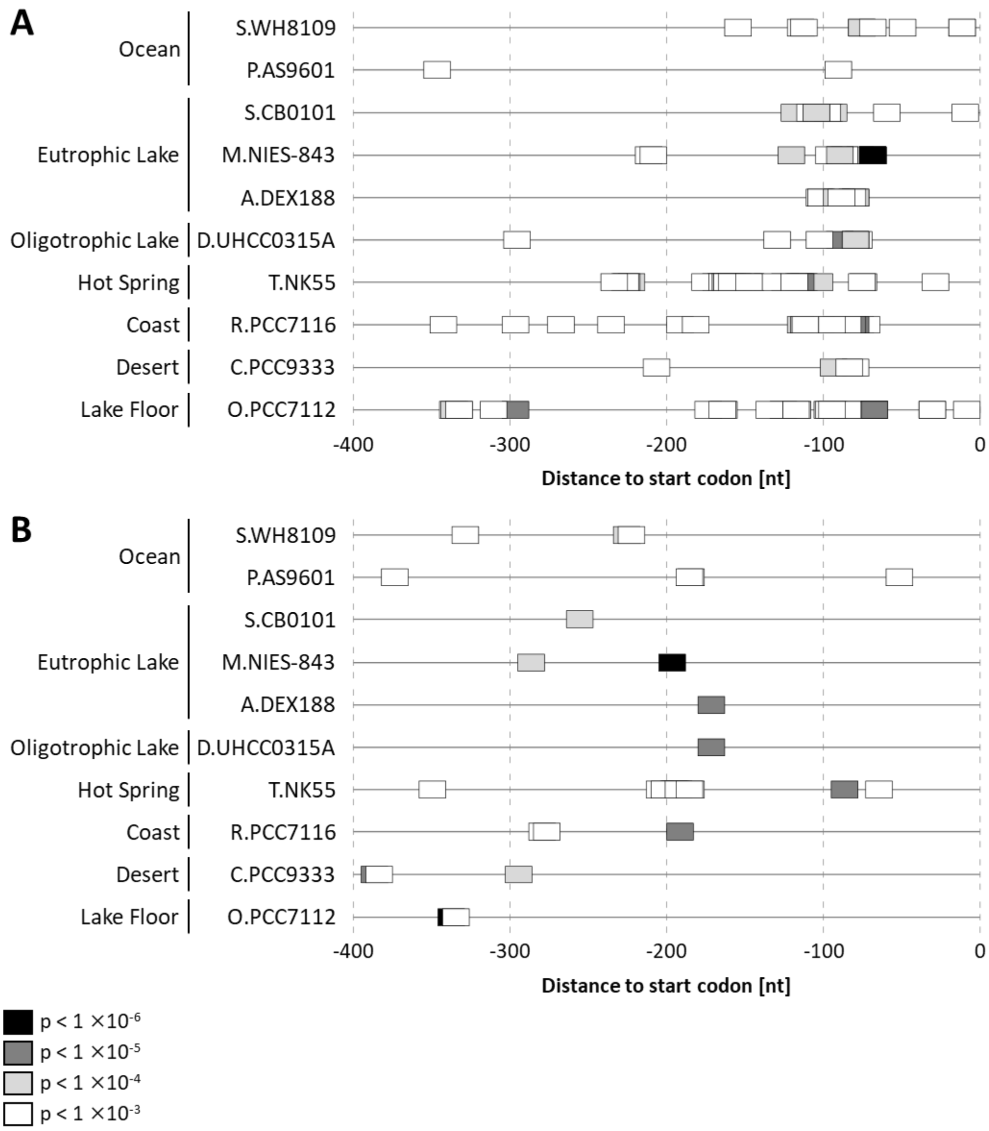


**Figure S5.** The distribution of the HLR1 motifs in the upstream regions of *psbA* (A) and *psaA* (B) in each putative dominant species. The HLR1 motifs were searched within the 400-bp regions upstream of the start codons using FIMO in the MEME Suite.

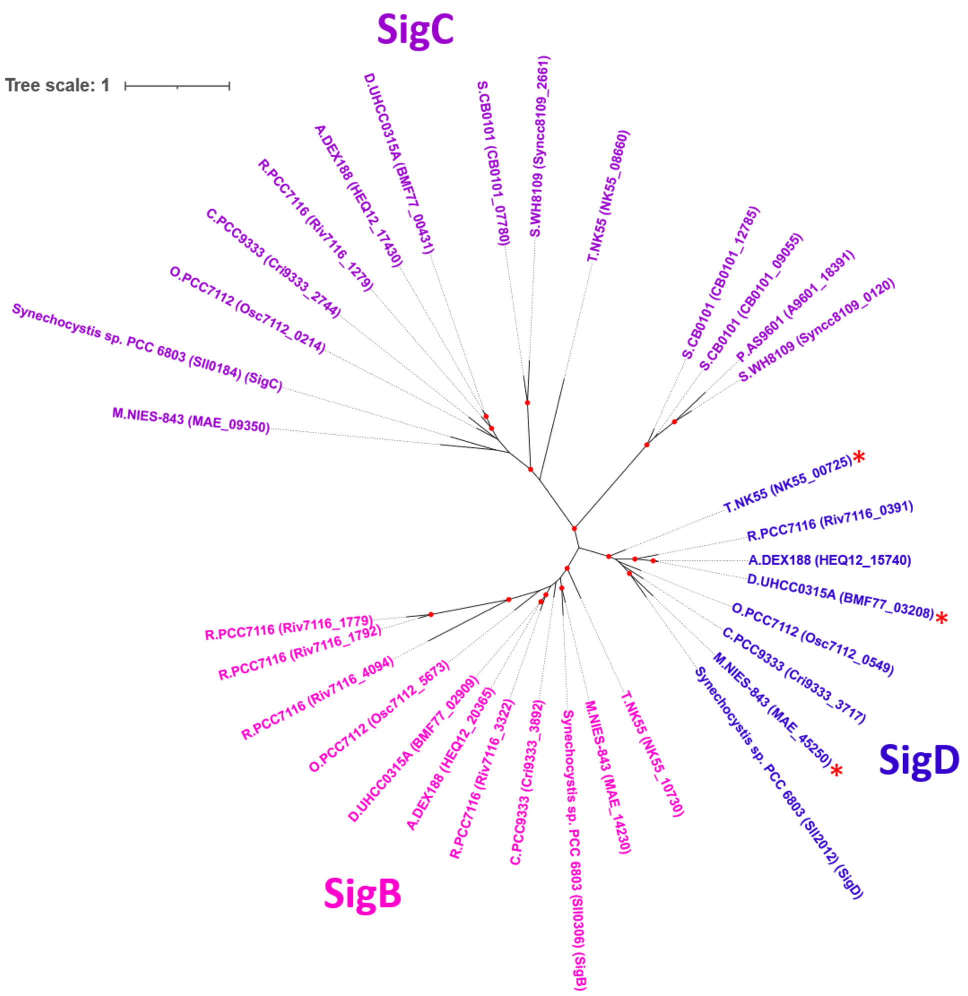


**Figure S6.** Maximum-likelihood phylogenetic tree of group 2 sigma factors SigB, SigC, and SigD assigned to orthogroup OG0000070 in the putative dominant species and *Synechocystis* sp. PCC 6803. Red asterisks indicate proteins encoded by the *psbA*-co-expressed genes (Table S7). Red dots indicate nodes with ultrafast bootstrap support values of ≥95.

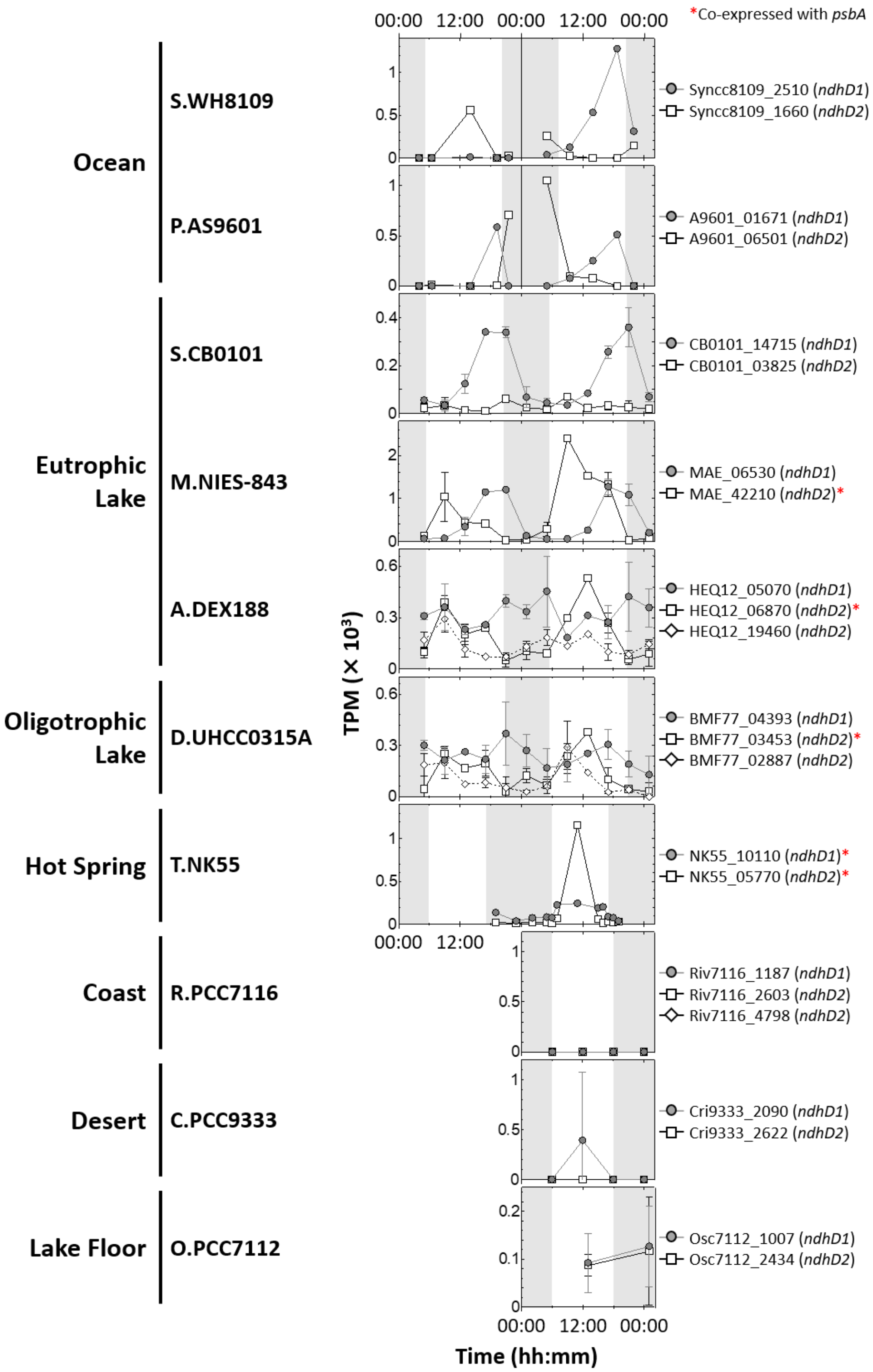


**Figure S7.** Diurnal expression patterns of *ndhD1* and *ndhD2* assigned to orthogroup OG0000120. In each plot, white and gray backgrounds indicate periods with and without sunlight, respectively. Red asterisks indicate genes included in the *psbA*-co-expressed gene sets (Table S7).

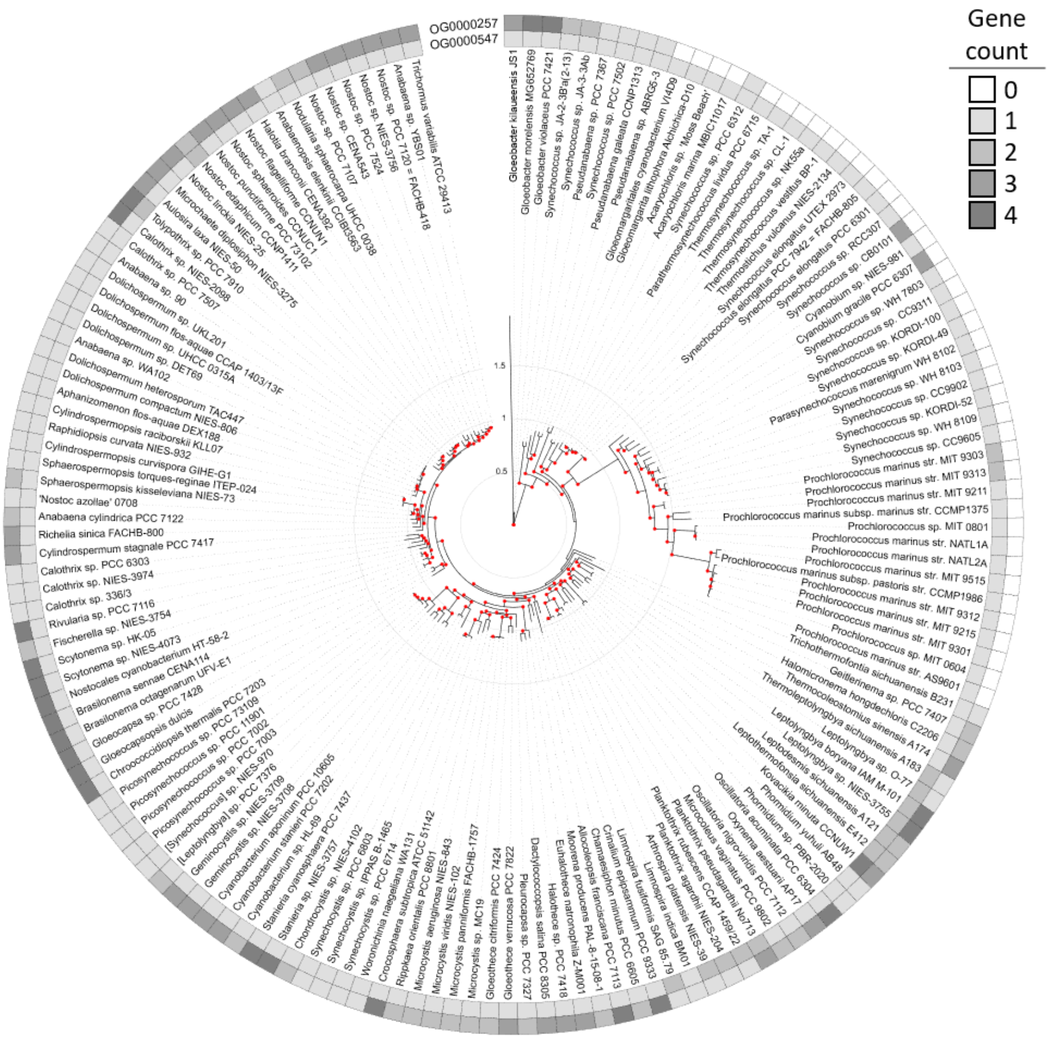


**Figure S8.** Heatmap showing the copy numbers of genes assigned to orthogroups OG0000547 and OG0000257, displayed around the phylogenetic tree of 159 cyanobacterial strains. The phylogenetic tree was inferred using the maximum-likelihood method based on a concatenated alignment of 126 single-copy orthologs. The scale bar represents evolutionary distance in amino acid substitutions per site. Red dots indicate nodes with ultrafast bootstrap support values of ≥95.

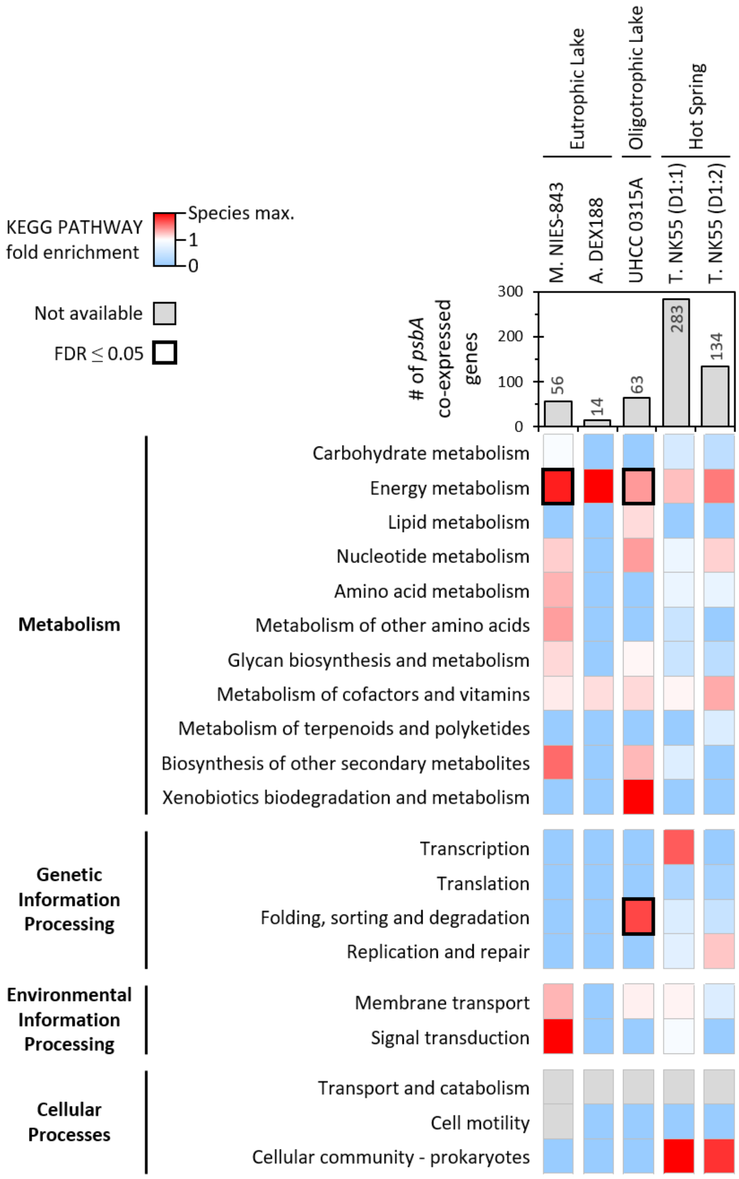


**Figure S9.** Heatmap showing fold enrichment of KEGG PATHWAY categories among genes co-expressed with *psbA*. Entries with FDR < 0.05 are outlined with thick borders.

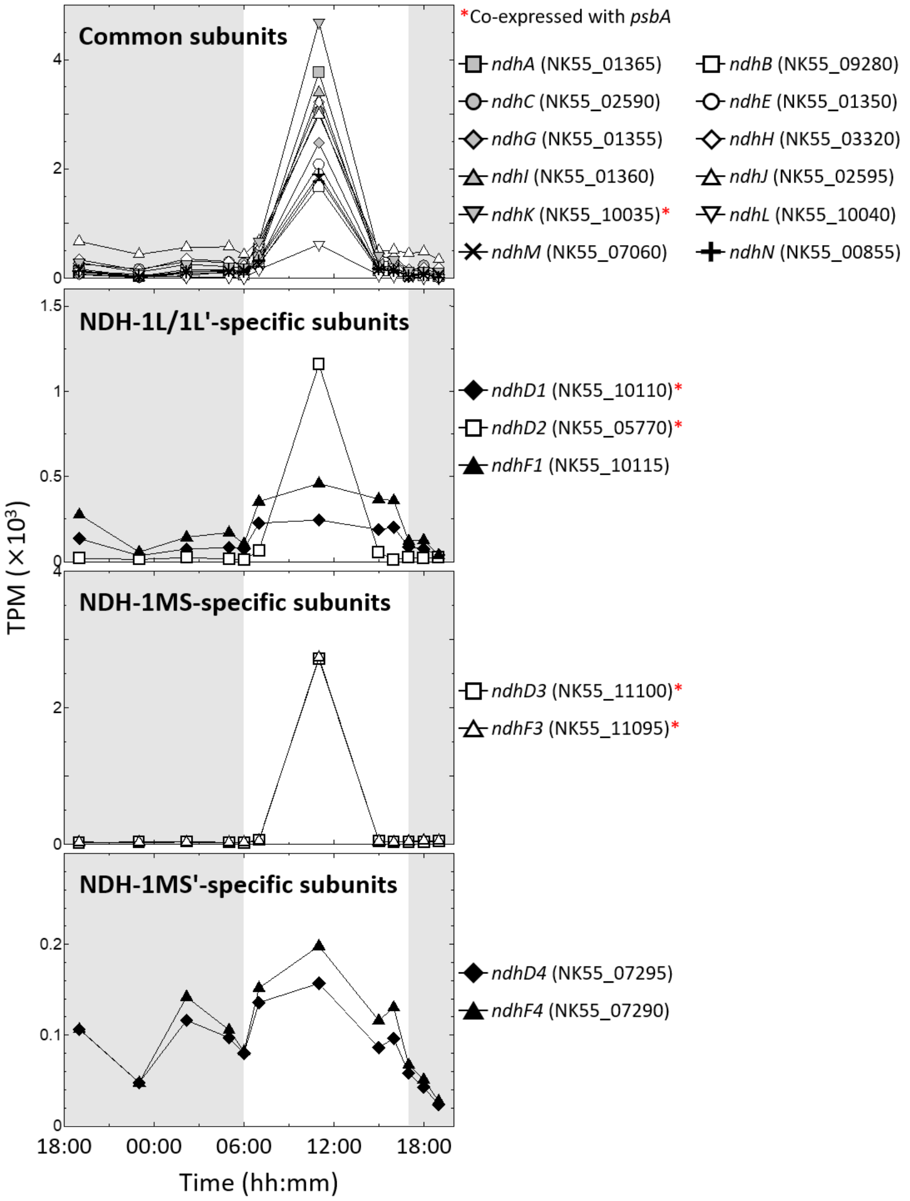


**Figure S10.** Diurnal expression patterns of genes encoding NDH-1 complex subunits in T.NK55 from the Hot Spring dataset. In each plot, white and gray backgrounds indicate periods with and without sunlight, respectively. Red asterisks indicate genes included in either the co-expressed gene set for the D1:1-encoding *psbA* or that for the D1:2-encoding *psbA* (Table S8).

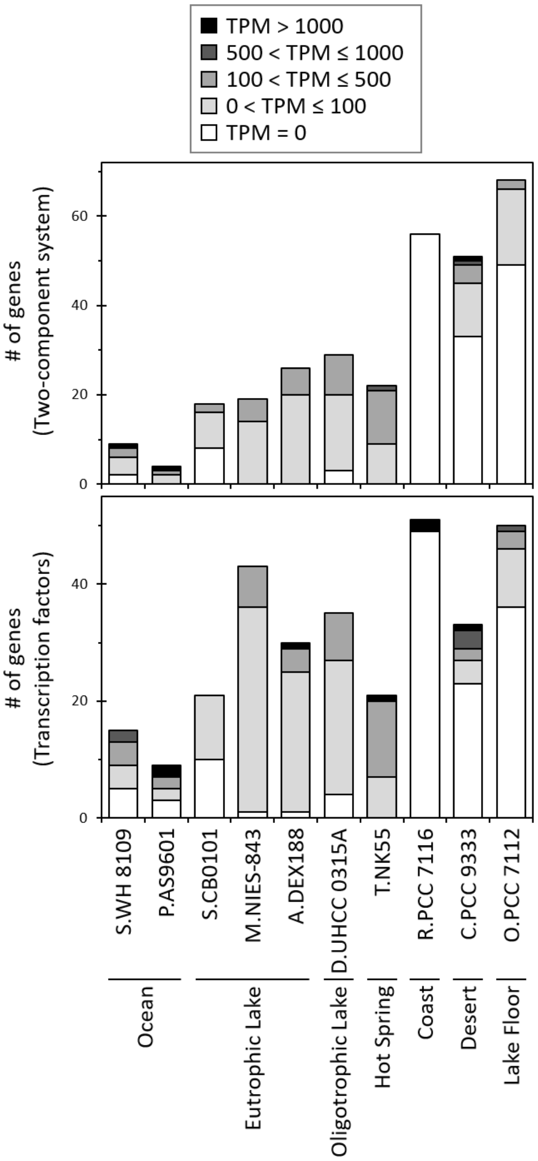


**Figure S11.** Stacked bar chart showing the total number of genes assigned to "Two-component system" and "Transcription factors" of the KEGG BRITE database and their TPM distributions. For each putative dominant species, the total number of genes in each category is divided into groups according to the TPM values of the individual genes.

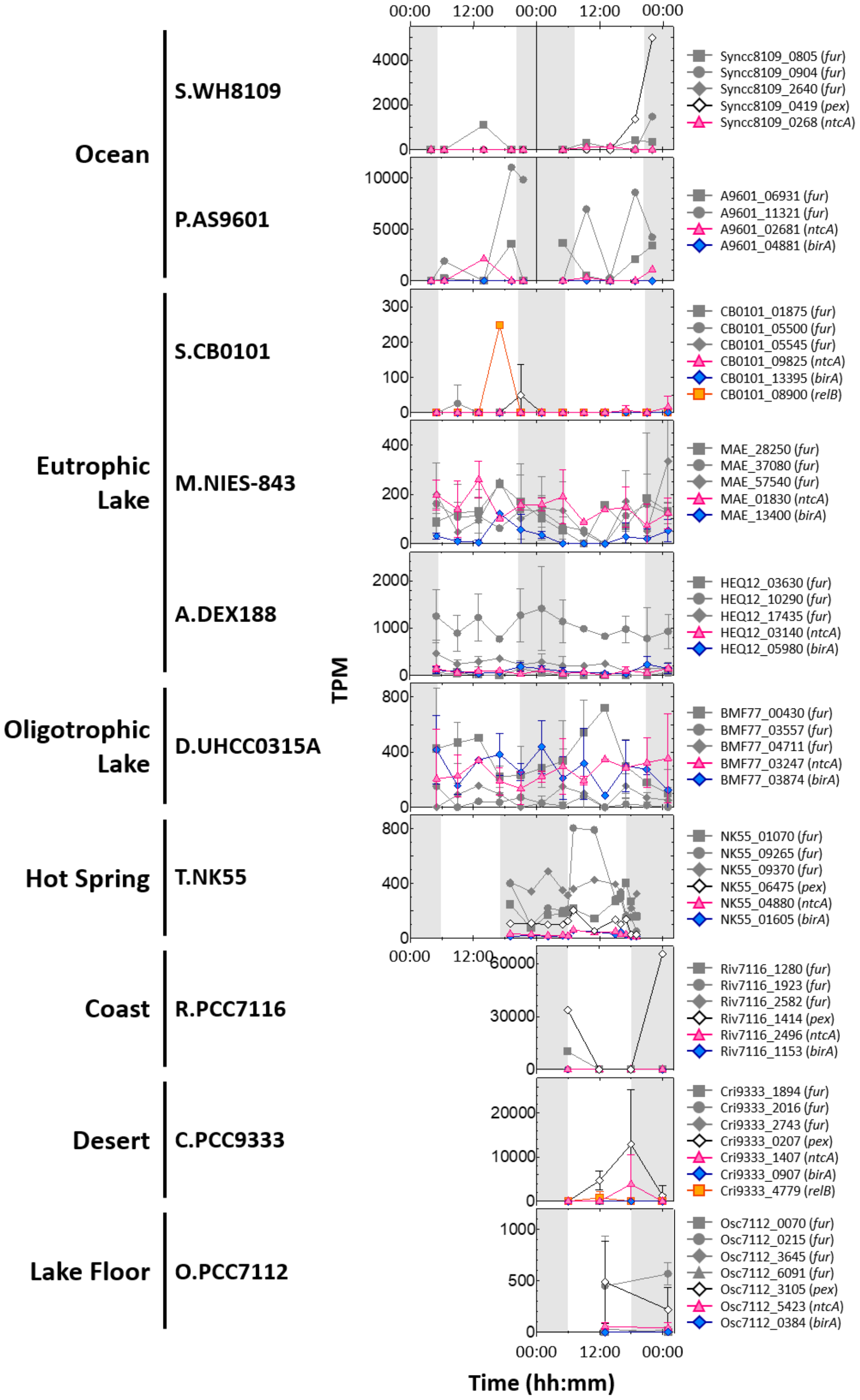


**Figure S12.** Diurnal expression patterns of genes assigned to "Transcription factors" of the KEGG BRITE database. In each plot, white and gray backgrounds indicate periods with and without sunlight, respectively. *fur* (K03711), gray symbols; *pex* (K26743), black open diamonds; *ntcA* (K21561), pink triangles; *birA* (K03524), blue diamonds; *relB* (K18918), yellow squares.

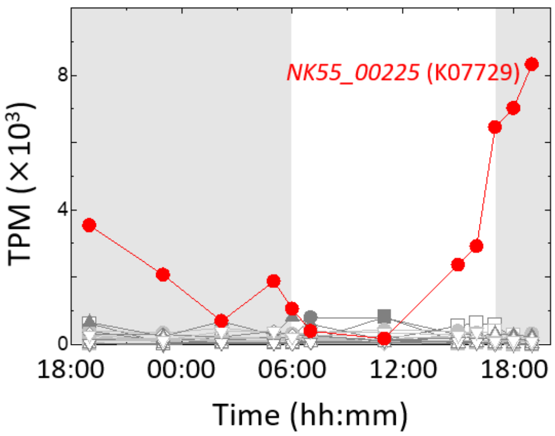


**Figure S13.** Diurnal expression patterns of all genes assigned to "Transcription factors" of the KEGG BRITE database in T.NK55 from the Hot Spring dataset. In each plot, white and gray backgrounds indicate periods with and without sunlight, respectively. NK55_00225, which is assigned KO identifier K07729, is shown with red symbols; all other genes are shown with gray symbols.
